# Connection “Stripes” in the Primate Insula

**DOI:** 10.1101/2020.11.03.361055

**Authors:** M. Krockenberger, T.O. Saleh, N.K. Logothetis, H.C. Evrard

**Affiliations:** Werner Reichardt Center for Integrative Neuroscience, Karl Eberhard University of Tübingen, Tübingen, Germany; Max Planck Institute for Biological Cybernetics, Tübingen, Germany; Centre for Imaging Sciences, Biomedical Imaging Institute, University of Manchester, Manchester, UK; Translational Neuroscience Laboratories Division, Center for Biomedical Imaging and Neuromodulation (C-BIN), Nathan Kline Institute for Psychiatric Research, Orangeburg, NY, USA

## Abstract

The insula has been classically divided into vast granular, dysgranular and agranular sectors. Over the years, several distinct studies proposed subdivisions of these sectors, with however no consensus. We recently proposed a cyto- and myelo-architectonic partition in which each sector contained sharply delimited areas (Evrard *et al.* 2014 J Comp Neurol 522: 64-97). Some of these areas were further divided into distinct subareas with obvious functional implications. Here, we examined the spatial relationship between architectonic boundaries and tract-tracing labeling in the insula in the macaque monkey. Injections of neuronal tracers in distinct areas of the prefrontal or anterior cingulate cortices produced heterogeneous and discontinuous patterns of anterograde and retrograde labeling in the insula. These patterns were made of sharply delimited patches forming anteroposterior stripes across consecutive coronal sections. While the overall pattern of labeling varied with the injection site, the patches systematically coincided with specific architectonic subareas, particularly in the dysgranular insula. This unequivocally validates our prior architectonic partition and strongly supports the idea of a refined modular *Bauplan* of the primate insula. This modular organization may underlie a serial stream of integration of interoception with ‘self-agency’ and ‘social’ activities across distinct insulo-prefrontal processing units that need to be explored.

## Introduction

The insular cortex constitutes the fifth lobe of the cerebral cortex in anthropoid primate (Reil 1809). Long perceived as a rather primitive and poorly organized limbic cortex, it recently re-emerged as having a crucial role in representing the physiological state of the organs and tissues of the body (Craig 2002), and in contributing to emotional embodiment, with a role in shaping cognitive processes including perhaps subjective perceptual awareness (Craig 2009; Gu *et al.* 2013; Salomon *et al.* 2016). These functions are of paramount importance in human mental health and both the insula is, together with the cingulate cortex, one of the major target common to several distinct neuropsychiatric disorders (Goodkind *et al.* 2015). Thus, understanding how the insula is anatomically and functional organized is critically needed.

Beyond the basic macro-morphological of gyri and sulci (Reil 1809; Naidich *et al.* 2004), the primary approach to assess the organization of a brain region is to examine its histological architecture (Brodmann 1909; Vogt and Vogt 1919). The insular cortex has been classically divided into vast cytoarchitectonic sectors with, over the years, various proposals for further partitioning (Fig. 1; Brodmann 1909; Vogt 1911; Rose 1928; Brockhaus 1940; Mesulam and Mufson 1982; Kurth, Eickhoff, *et al.* 2010; Gallay *et al.* 2012; Evrard *et al.* 2014). Brodmann (1909) divided the insula into a large anterior agranular sector and posterior granular sector (Fig. 1A). He however stated that further examination should be carried on. Recognizing a different orthogonal orientation, the Vogt’s (1911) recognized a dorsal granular sector and a ventral agranular sector that they further divided into myelo-architectonic areas aligned in some cases with the localization of the sulci (Fig. 1B). Rose (1928) and Brockhaus (1940) proposed much finer parcellations into multiple areas with rather abrupt cytoarchitectonic borders. They also added an intermediate dysgranular (or “mesocortical” with Brockhaus, and “propeagranularis” with Rose) sector in between the agranular and granular sectors already reported by Brodmann (Fig. 1C and D). Although Rose used only one human insula and did not represent the position of the sulci, taken together with the six insulae used by Brockhaus, their work suggested a parcellation into numerous (26 for Brockhaus to 31 for Rose) relatively sharply defined areas that readily suggested a compartmented functional organization. Later in the century, after a resurgence of interest for the insular cortex driven by new tract-tracing, lesion and recording studies primarily in non-human primates, a series of authors opted for a simpler parcellation recognizing only the three original sectors (Roberts and Akert 1963; Sanides 1968; Burton and Jones 1976; Mesulam and Mufson 1982) (*e.g.*, Fig. 1F). In their seminal tract-tracing studies in the macaque monkey, Mesulam and Mufson divided the insula into two vast territories, with a vague gradual separation crossing the middle of the dysgranular sector, with no recognition of a finer internal organization (Fig. 1G) (Mesulam and Mufson 1982; Mufson and Mesulam 1982; Mesulam and Mufson 1985; Mufson *et al.* 1997). The interest for the insula sparked again with new findings in functional imaging in humans and with the proposal of a crucial role of the insula interoception (Craig 2002). Using perikaryal stain with a well-established “observer-independent” and probabilistic parcellation method (Amunts *et al.* 2002), Kurth and colleagues map the human insula posterior to its central sulcus (Kurth, Eickhoff, *et al.* 2010). They proposed a consistent division of the granular and dysgranular sectors into two and three sharply-delimited areas, respectively (Fig. 1E). Using multiple histological and immunohistochemical stains in macaque monkeys and humans, Morel, Gallay and colleagues also recognized a finer division of the classical sectors (*e.g.*, Fig. 1H) (Gallay *et al.* 2012; Morel *et al.* 2013). However, they did not observe any consistent coincidence between the borders defined by the different stains. Finally, using cyto- and myelo-architectonics in 20 cynomolgus macaque monkeys, Evrard and collaborators reported the parcellation of the classical granular, dysgranular and granular sectors into respectively 4, 4 and 7 sharply delimited and highly consistent areas (Evrard *et al.* 2014). Three dysgranular areas and one agranular area were further subdivided into subtle sub-areas with optimal coincidence of the boundaries defined by the cyto- and myelo-architectures. This new map was relatively consistent with the map of the posterior insula by Kurth et al. (2010) and with map of the anterior agranular insula, anterior to the limen, by Carmichael and Price (1994) where cross-stain coincidence was also reported. (For an exhaustive comparison of these different maps, see the *Discussion* in Evrard *et al.*, 2014.)

**Figure 1.**
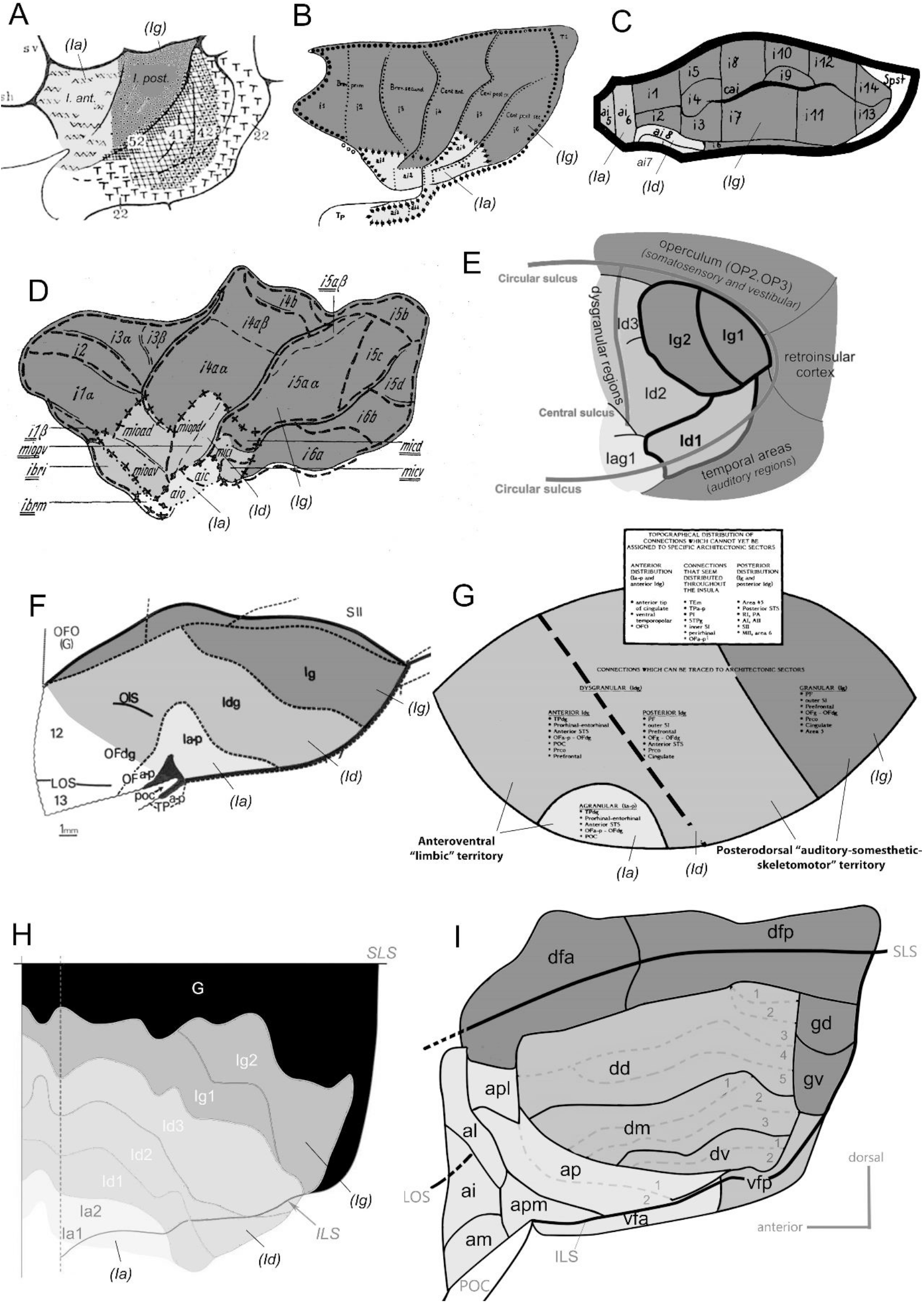
Flat maps drawings of the insular cortex showing the delineation of distinct sectors, areas or sub-areas in seven different cyto- and/or myelo-architectonic studies in humans or cercopithecine monkeys. **A.** Cytoarchitectonic division of the human insula into one anterior agranular (*I. ant.*; *Ia*) and one posterior granular (*I. post.*; *Ig*) sector (adapted with permission from Brodmann 1909). **B.** Myeloarchitectonic division of the human insula in which six myeloarchitectonic fields (*i1-i6*) were also cytoarchitectonically isocortical (or granular; *Ig*) and seven others (*ai1-i7*) allocortical (or agranular; *Ia*) (adapted with permission from Vogt 1911). **C.** Cytoarchitectonic division of the hamadryas baboon (*Papio hamadryas*) insula into two agranular (ai7 and ai8; *Ia*), two dysgranular (ai5 and ai6; *Id*), and fourteen granular areas (i1-14; *Ig*) (adapted from Rose 1928). Rose’s agranular and dysgranular sectors contain two (ai1 and ai10) and seven (ai2-6, ai9) additional agranular and dysgranular areas, respectively, that are not shown in this figure. **D.** Cyto- and myelo-architectonic division of the human insula into two allocortical (agranular; *Ia*), eight mesocortical (dysgranular; *Id*) and 16 isocortical (granular; *Ig*) areas (adapted from Brockhaus 1940). **E.** Cytoarchitectonic division of the human insula posterior to central sulcus, with a subdivision of the granular and dysgranular insula into two and three areas, respectively (adapted with permission from Kurth, Eickhoff, *et al.* 2010). **F**. Cytoarchitectonic division of the rhesus macaque (*Macaca mulata*) insula into one agranular (*Ia*), one dysgranular (*Id*) and one granular (*Ig*) sector (adapted from Mesulam and Mufson 1982). **G.** A similar view of the cytoarchitectonic map shown in panel E, overlaid with a hodological division into one anteroventral “limbic” territory and one posterodorsal “auditory-somesthetic-skeletomotor” territory (adapted with permission from Mesulam and Mufson 1982). The separation between the two territories crosses the middle of the dysgranular sector (thick dashed line in *Id*). **H.** Cytoarchitectonic division of the macaque insula into one vast “hypergranular” sector (G), two granular areas (Ig1 and 2), three dysgranular areas (Id1, 2 and 3) and two agranular areas (Ia1 and 2) (adapted with permission from Gallay *et al.* 2012). Note that the parcellation of the insula in the same subject (Mk4) but using other histological or immunohistochemical stains did not coincide with the cytoarchitectonic areas and with each other (see Gallay *et al.* 2012). **I.** Cyto- and myelo-architectonic subdivision of the three classical sectors of the macaque insula into four granular areas (*Idfa*, *Idfp*, *Igd*, *Igv*), four dysgranular areas (*Idd*, *Idm*, *Idv*, *Ivfp*), and seven agranular areas (*Ial*, *Iai*, *Iam*, *Iapl*, *Iap*, *Iapm*, *Ivfa*) (adapted from Evrard *et al.* 2014). Three of the dysgranular areas (*Idd*, *Idm* and *Idv*) and one of the agranular areas (*Iap*) were further subdivided in 2 to 5 subareas. In all panels, top is dorsal and left is anterior. The different degrees of cytoarchitectonic granularity are represented with different tones of gray (black, “hypergranular”; dark gray, granular; middle gray, dysgranular; light gray, agranular). For the abbreviations in A to G, see original publications. For the abbreviations in H, see the present abbreviations list. (adapted with permission from Evrard et al., 2014).

Cortical areas can be hard to define and identify, and their exact number in any species is uncertain (Kaas 2012). As expressed in our prior report, architectonic parcellation is biased by the subjective judgment of the observer and complicated by the inter-individual variability typically encountered across macaque brains (Evrard *et al.* 2014). In our architectonic examination, we consistently observed a thinner parcellation of the large dysgranular areas into narrow longitudinal sub-areas (Fig. 1H). However, these sub-areas were difficult to demonstrate to others in photomicrographs and were thus presented as “suggestions”. Neuronal tract-tracing adds an unbiased criterion for the parcellation of the cerebral cortex. The demonstration of an overlap between our cyto-/myelo-architectonic sub-areas and patterns of anterograde and/or retrograde labeling would provide an ideal demonstration of the validity of our insular parcellation. Therefore, in the present study, we examined the distribution of the labeling produced in the insula of the macaque monkey with injections of tracers in the prefrontal and cingulate cortices. We compared the distribution of the labeling with the architectonic parcellation defined in our prior study. We observed an optimal coincidence of architectonic and hodological boundaries.

## Material and Methods

### Tracer injection and histological processing

The present data was obtained from five adult rhesus macaque monkeys (*Macaca mulata*) of either sex. These monkeys were injected with neuronal tracers in the context of separate studies of the connectivity of the prefrontal cortex in the laboratory of Prof. Dr. Joseph L. Price at the Washington University of Saint-Louis, Missouri, USA. All animal protocols were approved by the Animal Studies Committee of Washington University and they complied with the guidelines of the NIH for the care and use of laboratory animals.

Detailed descriptions of the procedures for the microinjection of tracers, tissue fixation and histological processing have been previously published (*e.g.*, Carmichael and Price 1994; Ferry *et al.* 2000). Briefly, the tracer injections were made under deep general anesthesia and aseptic surgical conditions using glass micropipettes positioned through craniotomies at specific stereotaxic coordinates to reach selected cortical targets. The anesthesia was induced with ketamine (10 mg/kg, i.m.) and xyalzine (0.67 mg/kg, i.m.) and maintained with a gaseous inhalation of a mixture of oxygen, nitrous oxide, and halothane. A long-lasting analgesic (buprenorphine; 0.1 mg/kg, i.m.) was injected after the surgery as the animal recovered from the anesthesia. The tracers included biotinylated dextran amine (BDA), lucifer yellow (LY) and fluororuby (FR), at 5% or 10% in a 0.001 M phosphate buffered saline (pH 7.4) (PBS). The volumes injected varied between 0.3 and 1.2 μL.

After a survival period of ~14 days, the animals were sedated with ketamine (10 mg/kg, i.m.), deeply anesthetized with sodium pentobarbital (30 mg/kg, i.p. or i.m.) and transcardially perfused with PBS, followed by a sequence of 4% paraformaldehyde fixative phosphate buffers (1 liter of 4% paraformaldehyde, pH 6.5, 2 liters of 4% paraformaldehyde and 0.05% glutaraldehyde, pH 9.5, and finally 1 liter of the same solution with 10% sucrose). The brains were removed, blocked in the coronal plane and post-fixed in a 30% sucrose PBS at 4°C. After 2 to 6 days, the blocks of the brain were frozen and alternating 1-in-10 series of 50-μm-thick coronal sections were made using a sliding microtome, collected in PBS, and processed for histology or immunohistochemistry. BDA, LY and FR were revealed in separate series using the avidin-biotin-peroxidase staining method (Carmichael and Price 1995). One additional series was stained with the Nissl method for neuronal cell bodies and, in two cases, one series was stained with the Gallyas method for myelin (Carmichael and Price 1994).

### Data analysis

In a first analysis session, the location of the injection site in the prefrontal or cingulate cortex, and of the anterograde and/or retrograde labeling in all serial sections of the insular cortex ipsilateral to the injection site were examined at 5x, 10x, 20x and/or 40x using an upright brightfield microscope (Olympus BH2) and manually plotted using optical encoders attached to the microscope stage and interfaced with a personal computer (Minnesota Datametrics, St. Paul, MN, USA). This plotting system allowed a precise mapping of the location of retrogradely labeled cell bodies and an approximate, general representation of the location and density of the anterograde labeling. Composite of consecutive coronal maps of the labeling overlaid on maps of structural landmarks (cortical gray matter, delineation of cortical granular layer 4, claustrum…) were assembled in Adobe Photoshop (San Jose, CA, USA) and used to generate flat map representations (see below) of the overall distribution of the anterograde and retrograde labeling in the insula.

In a second analysis session, one to four representative series of coronal sections of the insula were selected in each case for a more precise mapping of the anterograde (and retrograde) labeling. These sections were photomicrographed at high resolution (0.2 × 0.2 × 1 microns, 21 z-stack levels) using the 10x and 20x planapochromatic objectives of a motorized microscope slide scanner (AxioScan.Z1, Carl Zeiss, Göttingen, Germany). The virtual sections were uploaded in a custom-made anatomical drawing software. The contour of the insula and its main anatomical landmarks (grey/white matter edge, layer 4, and claustrum) were drawn and the exact position of each retrogradely labeled neurons and anterogradely labeled axon terminal was plotted at high magnification while scrolling through the twenty-one 1-μm-tick focal points of the z-stack of the virtual sections to confirm the identity of the objects (i.e., stained cell body or axon terminal) (see *Results*).

In a third and independent analysis session, the cytoarchitecture and, when available, myeloarchitecture of the insula (Nissl- and Gallyas-stained sections) was examined by at least 2 independent examiners (MK, HCS and/or TOS) using both a stereomicroscope and the virtual sections. The areal and modular boundaries were drawn using architectonic criteria defined in prior studies (Carmichael and Price 1994; Evrard *et al.* 2014) (Fig. 1H). The plots of labeling and the corresponding architectonic maps were aligned together and figures were prepared in Adobe Photoshop (San Jose, CA, USA) without alteration other than to reach matching levels of brightness in all the photomicrographs. Finally, a lateral flat map view of the insula comparing the architecture and distribution of the anterograde/retrograde labeling was prepared for each case, using a procedure described in detail elsewhere (Evrard *et al.* 2014; see also *Results*).

## Results

### General observations

Figures 2A-E show drawings of the tracer injection sites in coronal sections from the five cases reported in this contribution (from A to E: OM85, OM42, OM27, OM30 and OM69). With the exception of OM85, these injection sites were described in previous articles (Ferry *et al.* 2000). All injections produced a dense core of tracer accumulation (black shapes in Fig. 2) and, in some cases, a diffuse halo of tracer diffusion surrounding the core (dark gray shapes in Fig. 2). The injection of LY in OM85 spread across all cortical layers within a small portion of the cingulate area 24b’ (Fig. 2A). The injection of BDA in OM42 reached the medial portion of the orbital prefrontal area 13l with a negligible spread to area 13m (Fig. 2B) (Ferry *et al.* 2000). The injection of BDA in OM27 was confined to a small portion of the orbital prefrontal area 11l with a thin halo spreading along the pipette track in area 46 (Fig. 2C) (An *et al.* 1998). The injection of BDA in OM30 filled a rather large portion of the orbital prefrontal area 12m with a small spread of the halo to area 12l (Fig. 2D) (An *et al.* 1998). The injection of LY in OM69 filled a rather large portion of the lateral prefrontal area 12r (Fig. 2E) and spread posteriorly in small portions of the dorsolateral prefrontal areas 45 and 46 (not shown) (Saleem *et al.* 2008).

**Figure 2.**
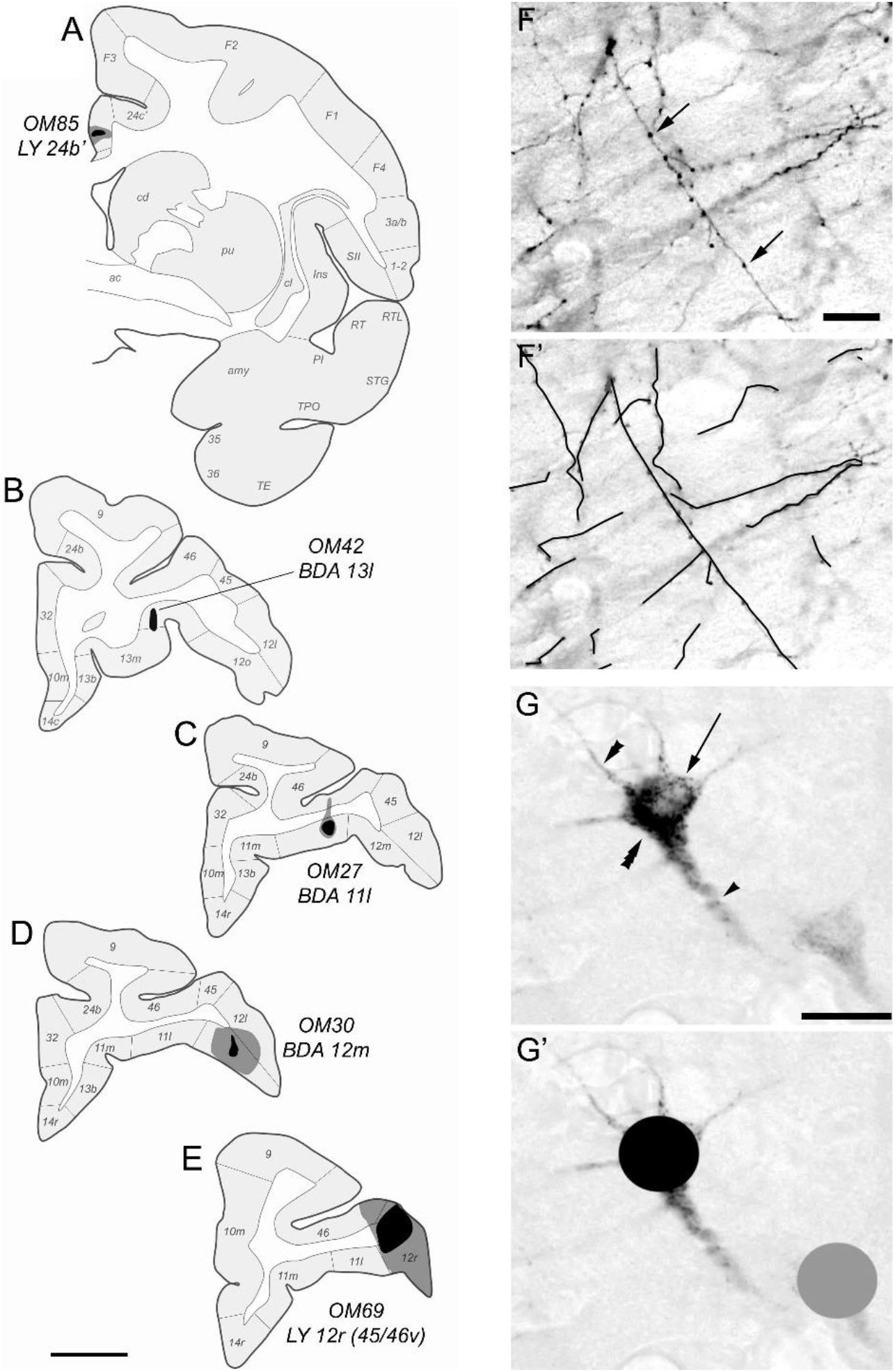
**A-E.** Drawings of the sites of injection of neuronal tracers in coronal sections of the right hemisphere of the cerebral cortex in five rhesus macaque monkeys. Each injection site is represented by a black core surrounded, in some cases, by a gray halo of diffusion. The five different cases are laid out from the most posterior (OM85) to the most anterior (OM69) injection. **A.** Injection site of lucifer yellow (LY) in area 24b’ in OM85. **B.** Injection site of biotin dextran amine (BDA) in area 13l in OM42. **C.** Injection site of BDA in area 11l in case OM27. **D.** Injection site of BDA in area 12m in case OM30. **E.** Injection of LY in area12r (and 45/46v, not shown) in OM69. Medial is left; dorsal is top. Scale bar = 5 mm. See the abbreviation list. **F.** Photomicrograph of representative retrogradely labeled pyramidal neurons in the insular cortex in case OM85 with an injection of lucifer yellow in area 24b’. The triple, double and single arrowheads point to the labeled perikarya, one of the basal dendrites and the apical dendrite, respectively. The arrow points to the unlabeled cell nucleus. **G.** Photomicrograph of representative anterogradely labeled axon terminals in the insular cortex in case OM85. The arrows point to examples of labeled varicosities localized along the axon fiber. **F’-G’.** Illustration of the labeling charting procedure. Scale bar = 10 μm.

All five injections produced anterograde and/or retrograde labeling in the insula. Figures 2F and G show high magnification photomicrographs of representative anterograde and retrograde labeling, respectively. Anterogradely labeled axon terminals were recognized by darkly stained cortico-cortical axonal fibers bearing varicosities (presumably synaptic boutons), identified as small beaded structures along the stained fibers (Fig. 2H). Each clearly distinct individual fiber bearing varicosities was charted using a continuous line (Fig. 2H’). The retrograde labeling occurred in typical cortico-cortical projection pyramidal neurons. It was identified by the staining of intracellular granules (presumably endocytoplasmic vesicles) filling the triangular soma and the proximal portion of the apical and basal dendrites while leaving the cell nucleus unstained (Fig. 2G). Intensely and weakly labeled perikarya (see the two neurons labeled in Fig. 2G) were charted in maps of the insula using black and gray symbols, respectively (Fig. 2G’).

Figures 3D to H show the flat map representations of the overall distribution of anterograde and/or retrograde labeling in the five cases. (Figures 3A-C illustrate the construction of these flat maps from series of consecutive coronal section drawings, using one section from OM85 as an example. See also Evrard *et al.* 2014.) The tracer injections made in distinct prefrontal or cingulate areas produced broadly distributed or rather restricted labeling, regardless of the size of the injection site. For example, the small injection of LY in area 24b’ in OM85 produced much more insular labeling (Fig. 3D) than the larger injection of BDA in area 12m in OM30 (Fig. 3G) or LY in area 12r in OM69 (Fig. 3H). In all cases but one (*i.e.*, OM30; Fig. 3G), the labeling occurred contiguously across several adjacent areas and sub-areas. For instance, in OM85, labeling was present in at least 11 consecutive architectonic entities throughout the dorso-ventral extent of the insula, with no ‘gap’ being entirely free of labeling (Fig. 3D). Yet, as illustrated in the figure and as will be demonstrated in the next section, the density (and to some extent layer distribution) of the labeling markedly and rather sharply varied throughout the insula, with a clear coincidence with the architectonic changes differentiating one cytoarchitectonic area or sub-area from the next.

**Figure 3.**
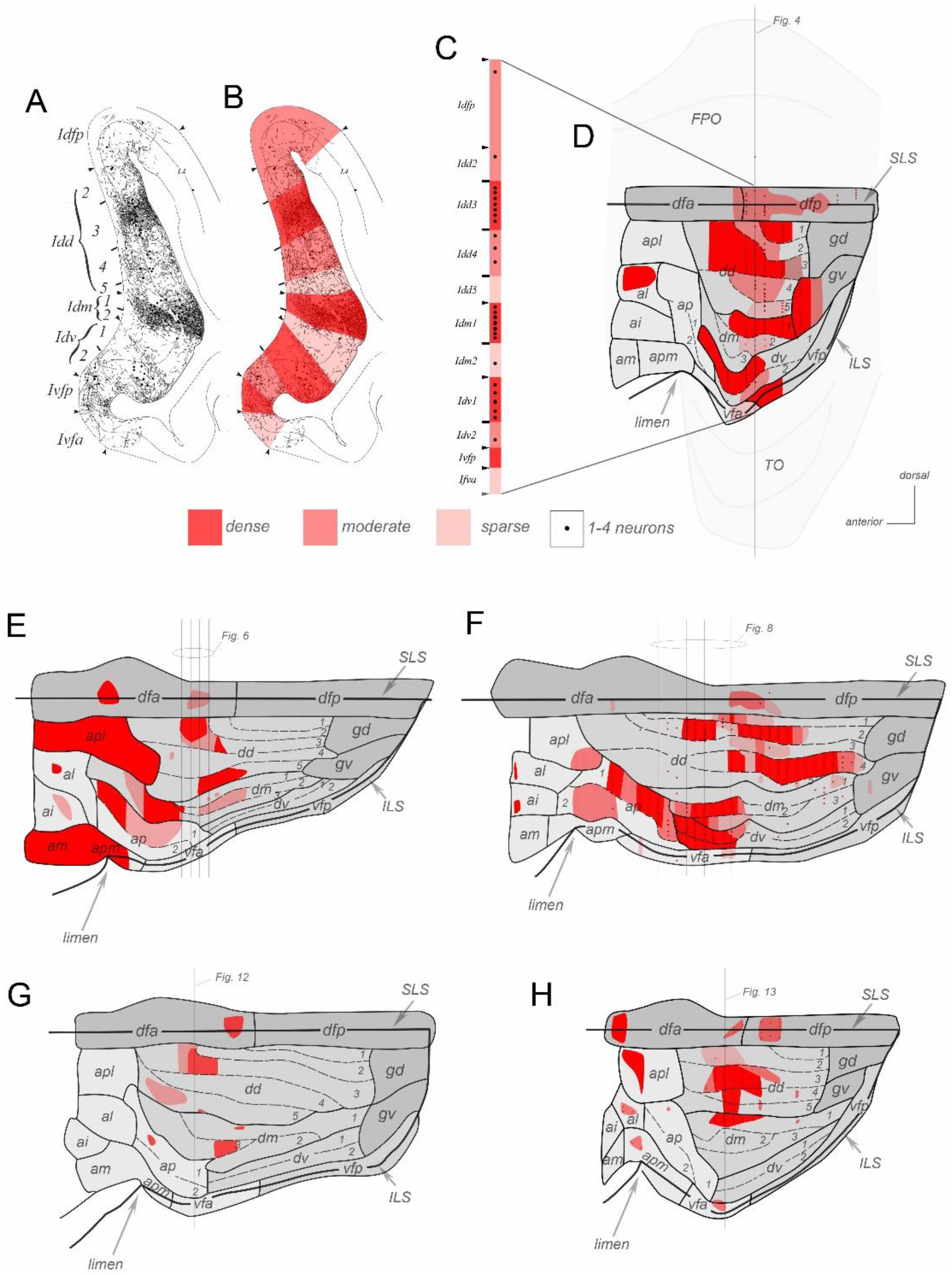
Flat map representations of the distribution of anterograde (shades of red) and retrograde (dark dots) labeling across the distinct architectonic areas and subareas of the insular cortex in five rhesus macaque monkeys with injections of tracers in different prefrontal or cingulate cortical areas. Panels A to C illustrate the method used to re-create the flat map of the insula shown in panel D for case OM85. The same principle was applied for all flat maps (Evrard *et al.* 2014). **A.** Original drawing of anterograde (thin lines) and retrograde (dots) labeling in one coronal section of the insular cortex in case OM85. **B.** Red-shades color-coding of three different densities of anterogradely labeled terminals (weak, moderate, dense). **C.** Transfer of the color-coding and overall number of retrogradely labeled neurons onto a flattened representation of the areas and labeling of the section. **D-H.** Flat map representations of the distribution of anterograde and/or retrograde labeling in the insula in cases OM85 (D), OM42 (E), OM27 (F), OM30 (G) and OM69 (H). In panel D, the frontoparietal (FPO) and temporal (TO) opercula are represented with a pale gray. In panels E-H, the representation of FPO and TO was omitted to avoid cluttering the figure. In all flat maps, the thick lines represent the superior and inferior limiting sulci (SLS and ILS). In all cases, the coronal sections were aligned using SLS, which is therefore represented by the straight line. The thinner plain contours delineate architectonic areas. The dashed contours delineate subareas within an area. The vertical lines crossing the maps indicate the anteroposterior level of the coronal sections shown in Figures 4 to 13 to illustrate the overlap of architectonic and hodological entities (see text).

### Architecto-hodological overlap

Figures 4 to 13 illustrate the overlap of tract-tracing labeling pattern variation with architectonic boundaries in representative coronal sections from each of the five cases. Figures 4, 6, 8, 12A-C and 13A-C show general low magnification views whereas Figures 5, 7, 9–11, 12D-E and 13E-F show corresponding high magnification demonstrations of the cyto- or myelo-architectonic boundaries. In all five cases, the pattern of anterograde and/or retrograde labeling was heterogeneous. The heterogeneity was characterized by isolated or multiple contiguous sharply delimited ‘patches’ of labeled cells and/or terminals. The metric and sharp delimitations of these patches were directly reminiscent of the architectonic sub-areas (or “modules”) defined in our prior contribution (Evrard *et al.* 2014). In fact, as detailed below, each patch of labeling overlapped indeed with a distinct architectonic entity. In general, the cyto- and myelo-architectonic features of the sub-areas described below for each case were comparable to the generic features reported in our earlier report (Evrard *et al.* 2014).

**Figure 4.**
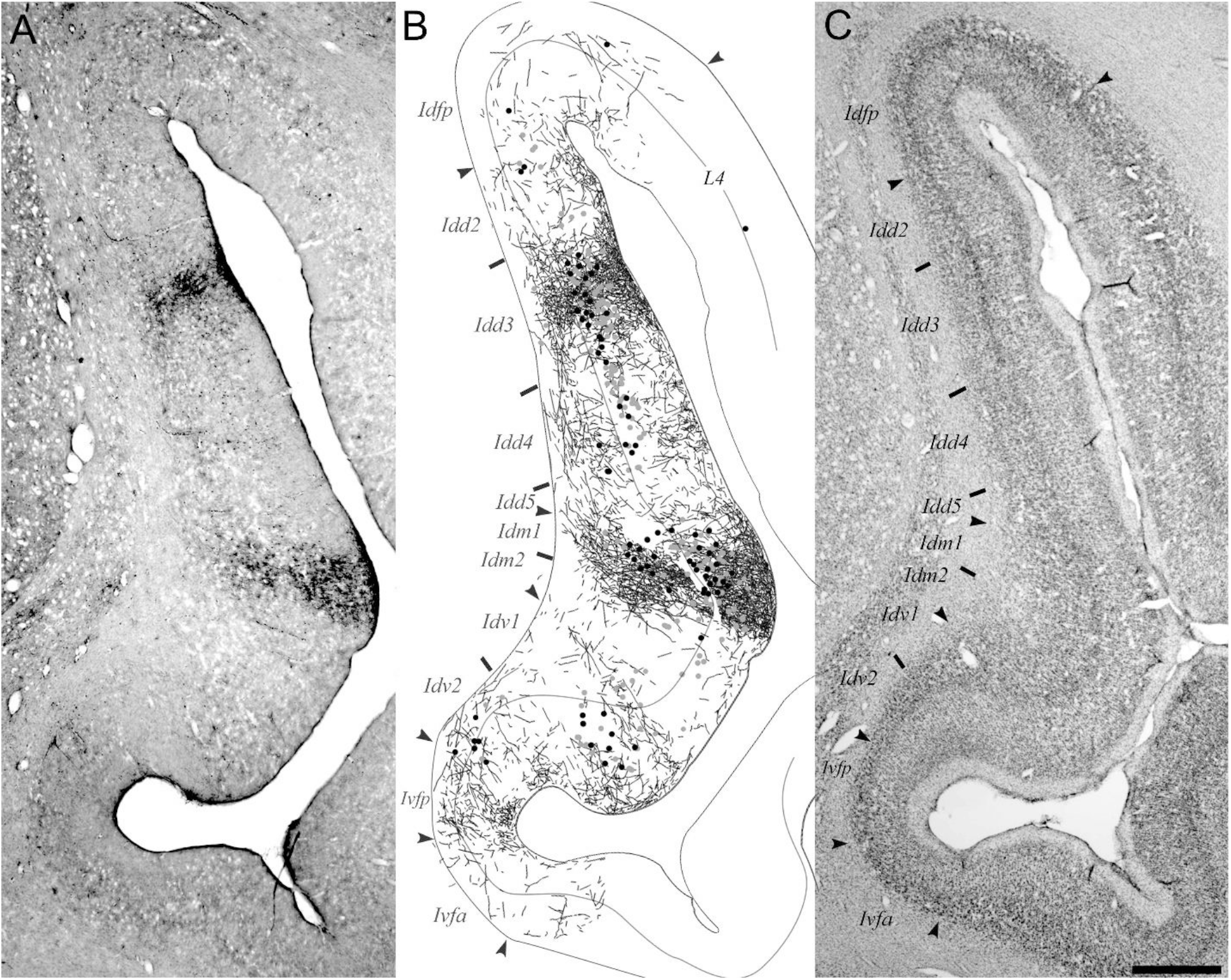
Composite showing the spatial overlap of patches of anterograde and retrograde labeling with specific cytoarchitectonic subareas of the insular cortex in case OM85 with an injection of lucifer yellow (LY) in cingulate area 24b’. **A.** Low magnification photomicrograph of a coronal section of the insula immunoreacted to visualize LY. Two patches of labeling are readily visible at low magnification. **B.** Plot of individual anterograde labeled terminals and retrogradely labeled perikarya made at high magnification in the same section as shown in panel A. Darkly and moderately stained perikarya are represented with black and gray dots, respectively. **C.** Low magnification photomicrograph of an adjacent Nissl-stained section showing the cytoarchitectonic boundaries between areas (arrowheads) and subareas (ticks). The same boundaries were transposed to the photomicrograph in panel B. Top is dorsal; left is medial. Scale = 1 mm.

**Figure 5.**
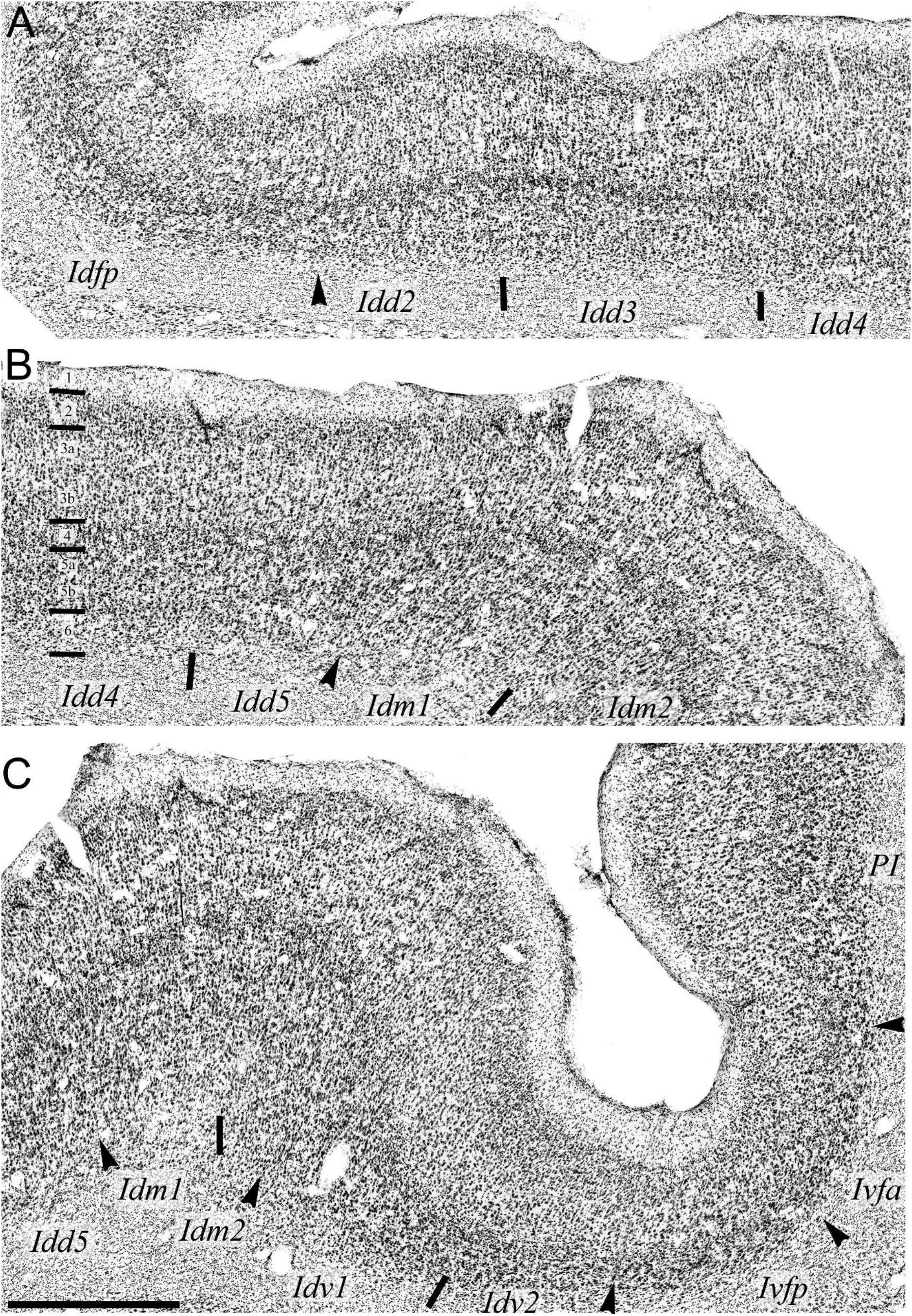
High magnification photomicrographs of the (A) dorsal, (B) middle and (C) ventral parts of the Nissl-stained section of the insula shown in panel C of Figure 4 in case OM85. The arrowheads and ticks mark the cytoarchitectonic boundaries between areas and subareas, respectively. The horizontal tick marks in panel B mark the separation of the cortical layers. Top is medial and left is dorsal. Scale bar = 1 mm.

**Figure 6.**
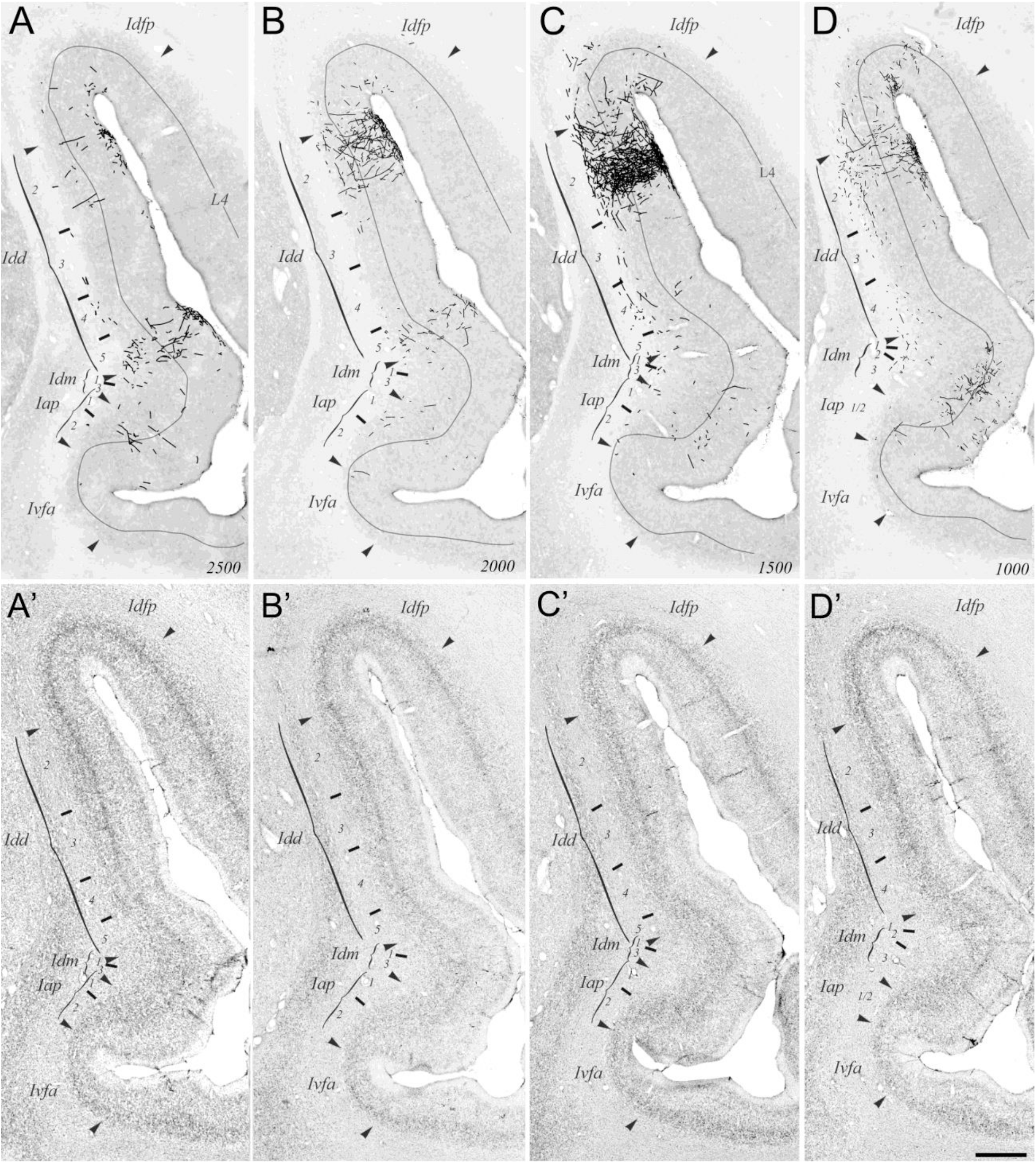
Composite showing the spatial overlap of patches of anterograde labeling with specific cytoarchitectonic subareas in the insular cortex in a case OM42 with an injection of biotin dextran amine (BDA) in the area 11l of the orbital prefrontal cortex (OPFC). **A-D.** Overlay of the charts of BDA-positive axon terminals on low-magnification photomicrographs of the corresponding BDA-stained sections of the insula across four coronal levels. The localization of the cytoarchitectonic boundaries between areas (arrowheads) and subareas (ticks) is shown for the subareas where anterograde labeling occurred. The distance of each coronal section from the level of the anterior commissure is noted in millimeters, in the bottom right corner of each panel (4500 mm in A to 0 mm in D). **A’-D’.** Overlay of the demarcation of the cytoarchitectonic boundaries of the insula on low-magnification photomicrographs of Nissl-stained sections directly adjacent to the BDA-stained sections shown in panels A-D. All areas and subareas are noted. (See abbreviations list.) High-magnification photomicrographs of selected subareas are shown in Figures 9, 10 and 11, allowing clear recognition of the cytoarchitectonic boundaries. (See text.) The asterisks mark the position of blood vessels used as landmarks across adjacent BDA- and Nissl-stained sections. Scale = 1 mm.

**Figure 7.**
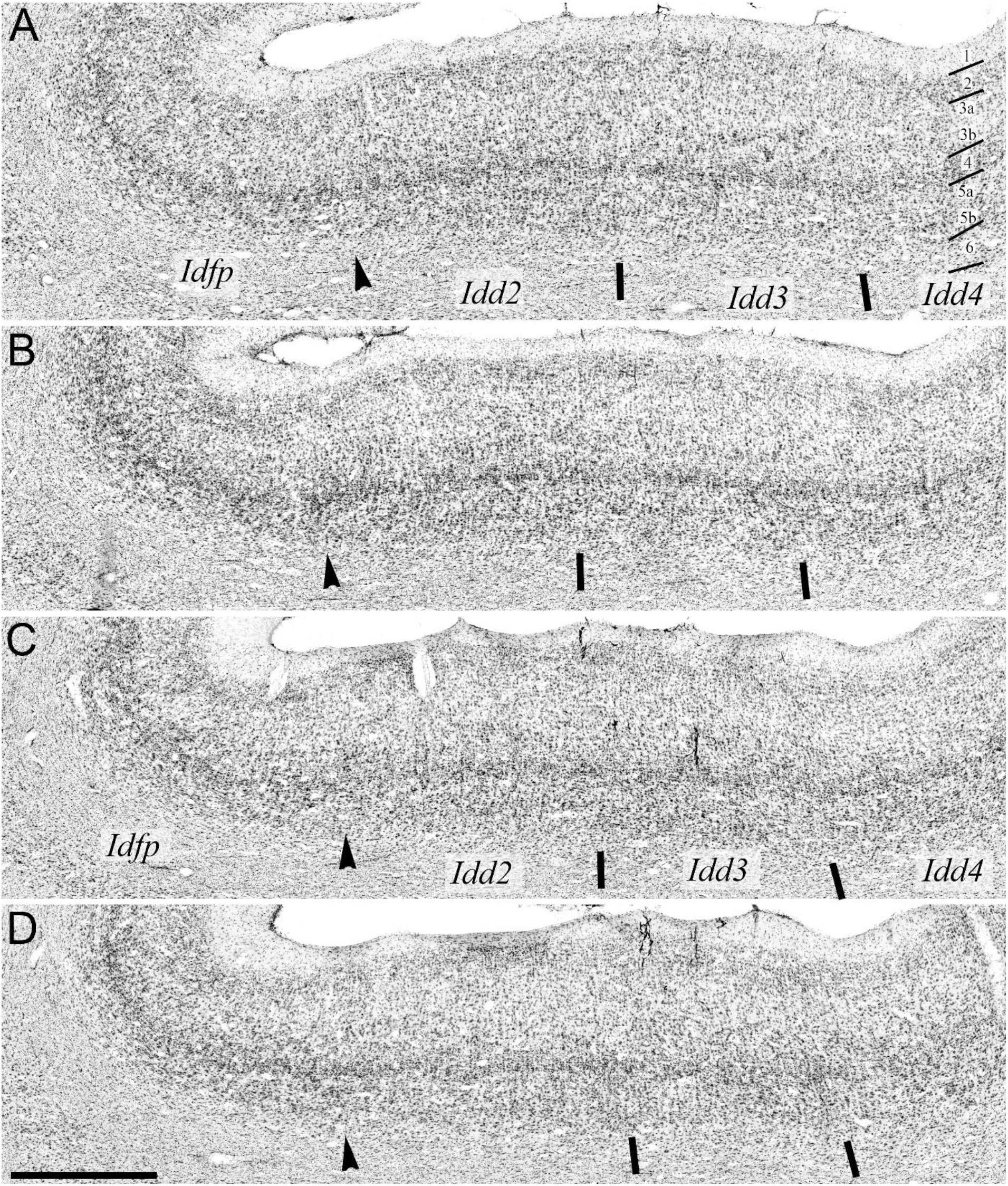
Overlay of the demarcation of cytoarchitectonic boundaries on high-magnification photomicrographs of four consecutive Nissl-stained sections, shown at low-magnification in Figure 6A’-D’ in case OM42. The emphasis is on Idfp, Idd2 and Idd3 where anterograde labeling occurred. Areas are separated by an arrowhead. Subareas are separated by tick marks. (See text and abbreviation list.) Scale = 1 mm.

**Figure 8.**
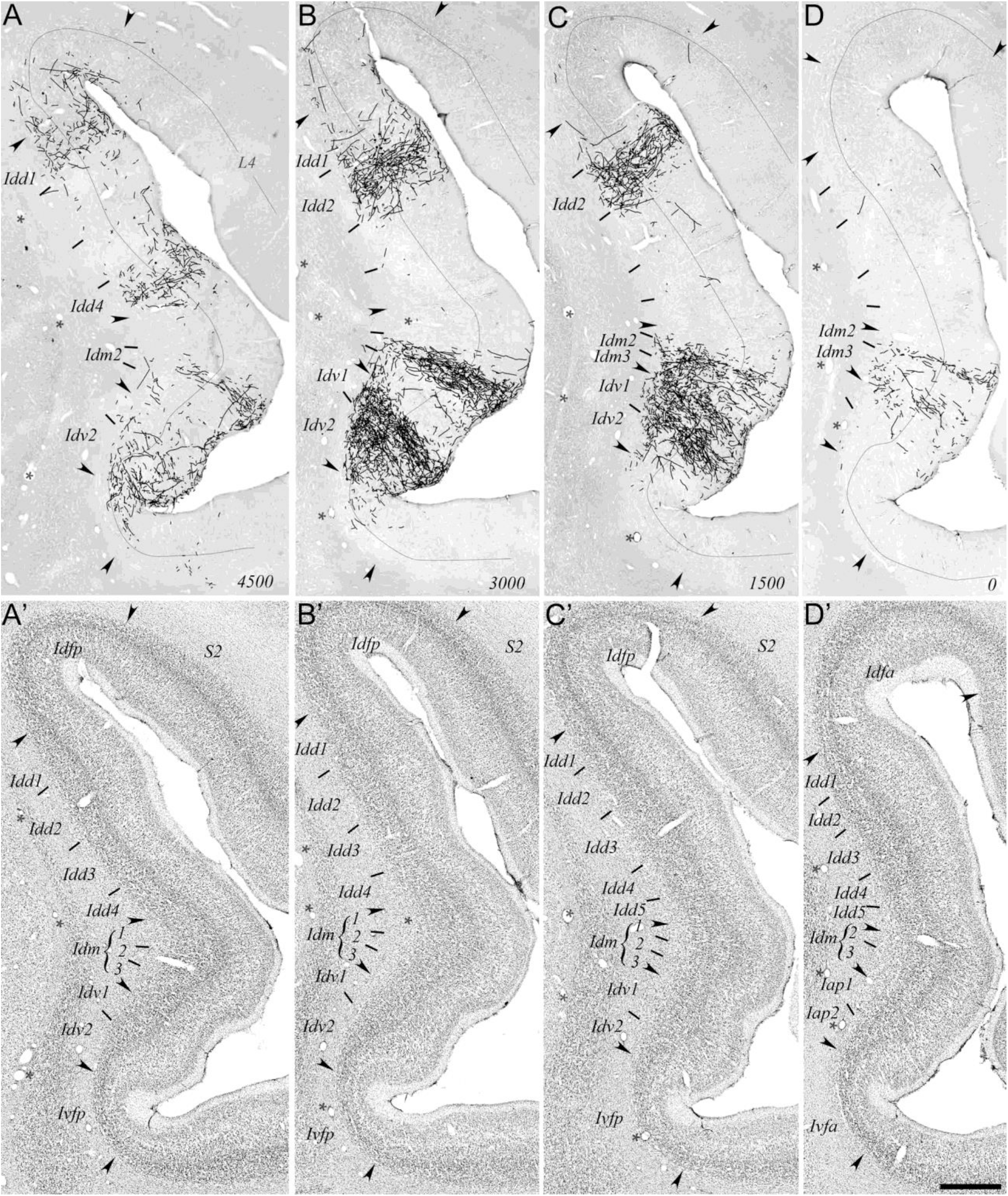
Composite showing the spatial overlap of patches of anterograde labeling with specific cytoarchitectonic subareas in the insular cortex in a macaque monkey (OM27) with an injection of biotin dextran amine (BDA) in the area 11l of the orbital prefrontal cortex (OPFC). **A-D.** Overlay of the charts of BDA-positive axon terminals on low-magnification photomicrographs of the corresponding BDA-stained sections of the insula across four coronal levels. The localization of the cytoarchitectonic boundaries between areas (arrowheads) and subareas (ticks) is shown for the subareas where anterograde labeling occurred. The distance of each coronal section from the level of the anterior commissure is noted in millimeters, in the bottom right corner of each panel (4500 mm in A to 0 mm in D). **A’-D’.** Overlay of the demarcation of the cytoarchitectonic boundaries of the insula on low-magnification photomicrographs of Nissl-stained sections directly adjacent to the BDA-stained sections shown in panels A-D. All areas and subareas are noted. (See abbreviations list.) High-magnification photomicrographs of selected subareas are shown in Figures 9, 10 and 11, allowing clear recognition of the cytoarchitectonic boundaries. (See text.) The asterisks mark the position of blood vessels used as landmarks across adjacent BDA- and Nissl-stained sections. Scale = 1 mm.

**Figure 9.**
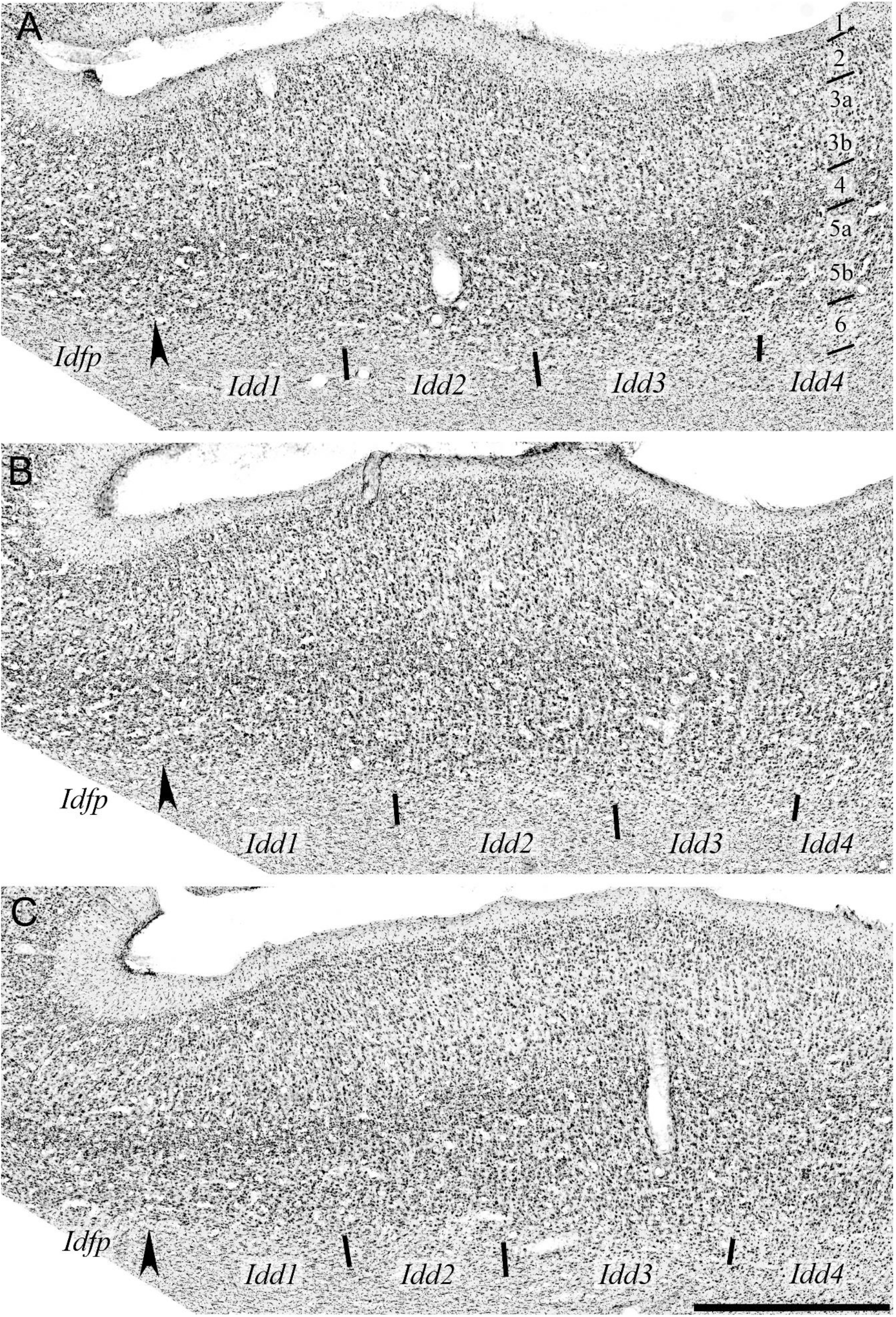
Overlay of the demarcation of cytoarchitectonic boundaries on high-magnification photomicrographs of three Nissl-stained sections, shown at low-magnification in Figure 8A’-C’ in case OM27. The emphasis is on subareas Idd1 and Idd2. Areas Idfp and Idd are separated by an arrowhead. Subareas 1, 2, 3 and 4 of Idd are separated by tick marks. (See text and abbreviation list.) Scale = 1 mm.

**Figure 10.**
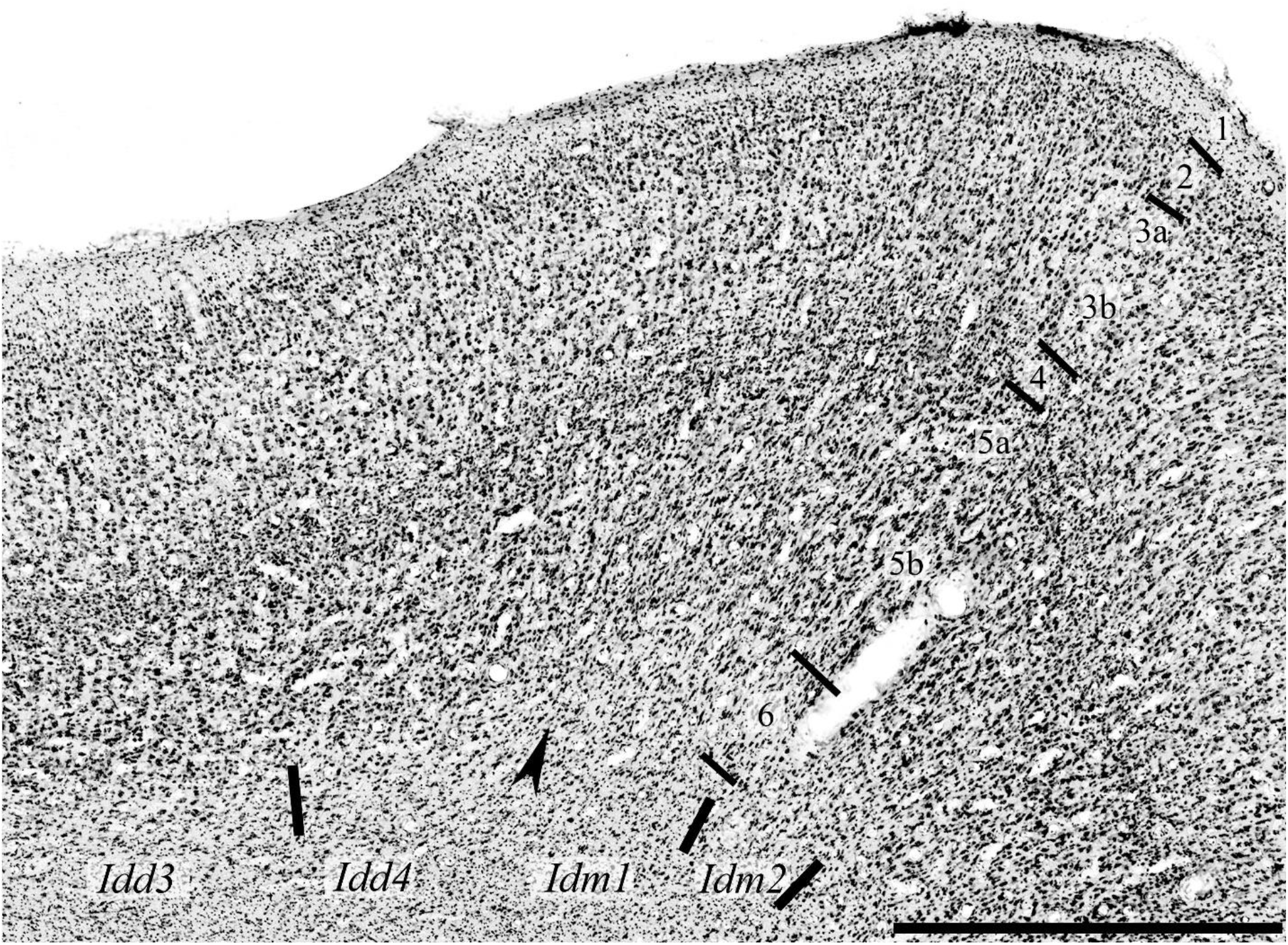
Overlay of the demarcation of cytoarchitectonic boundaries on high-magnification photomicrographs of one Nissl-stained section, shown at low-magnification in Figure 8A’ in case OM27. The emphasis is in on subareas Idd4. Subareas 3 and 4 of Idd are separated by tick marks. Areas Idd and Idm are separated by an arrowhead. (See text and abbreviation list.) Scale = 1 mm.

**Figure 11.**
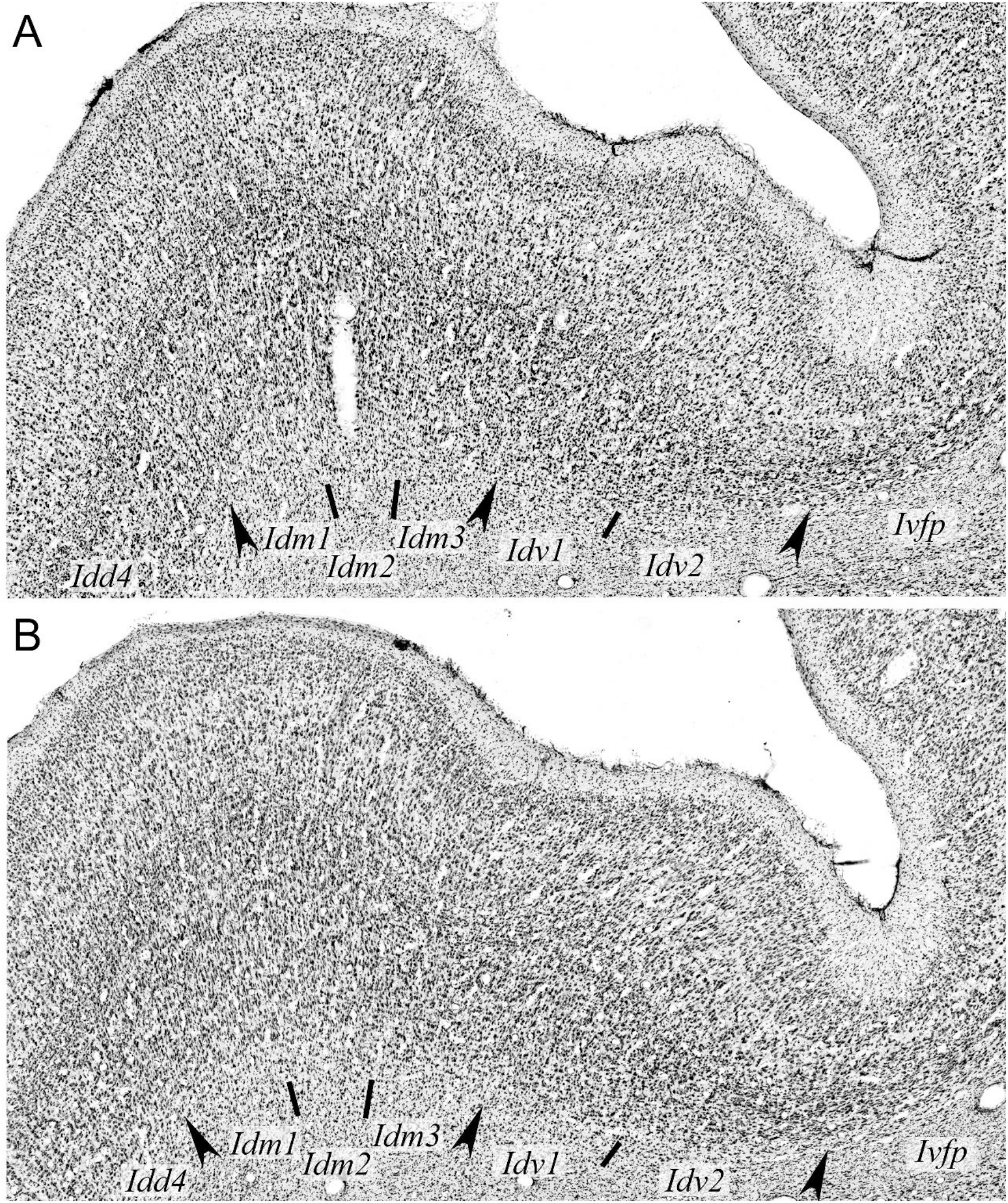
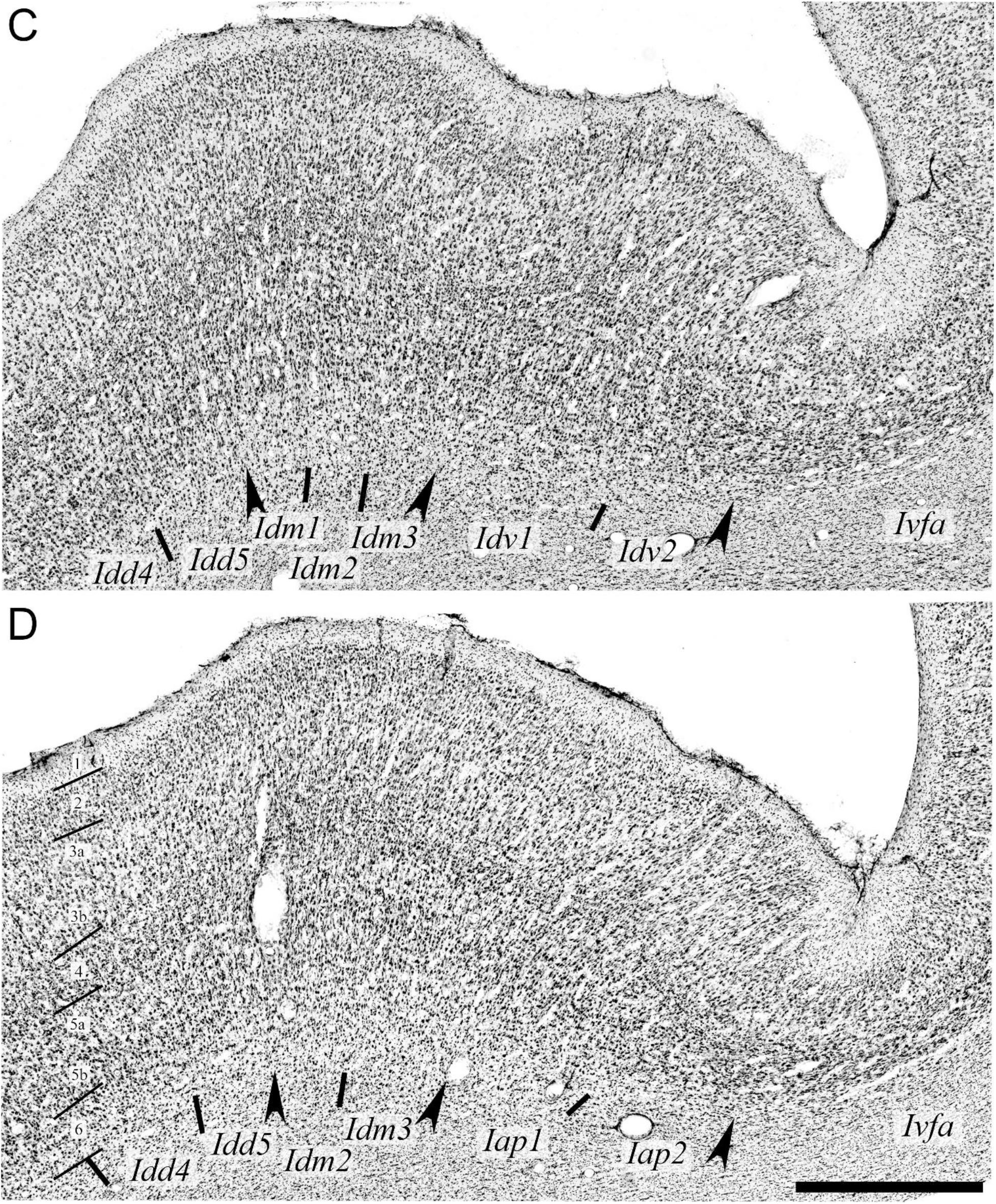
Overlay of the demarcation of cytoarchitectonic boundaries on high-magnification photomicrographs of four Nissl-stained sections, shown at low-magnification in Figure 8A’-D’ in case OM27. The emphasis is in on the subareas of areas Idm and Idv. Subareas are separated by tick marks. Areas are separated by arrowheads. (See text and abbreviation list.) Scale = 1 mm.

**Figure 12.**
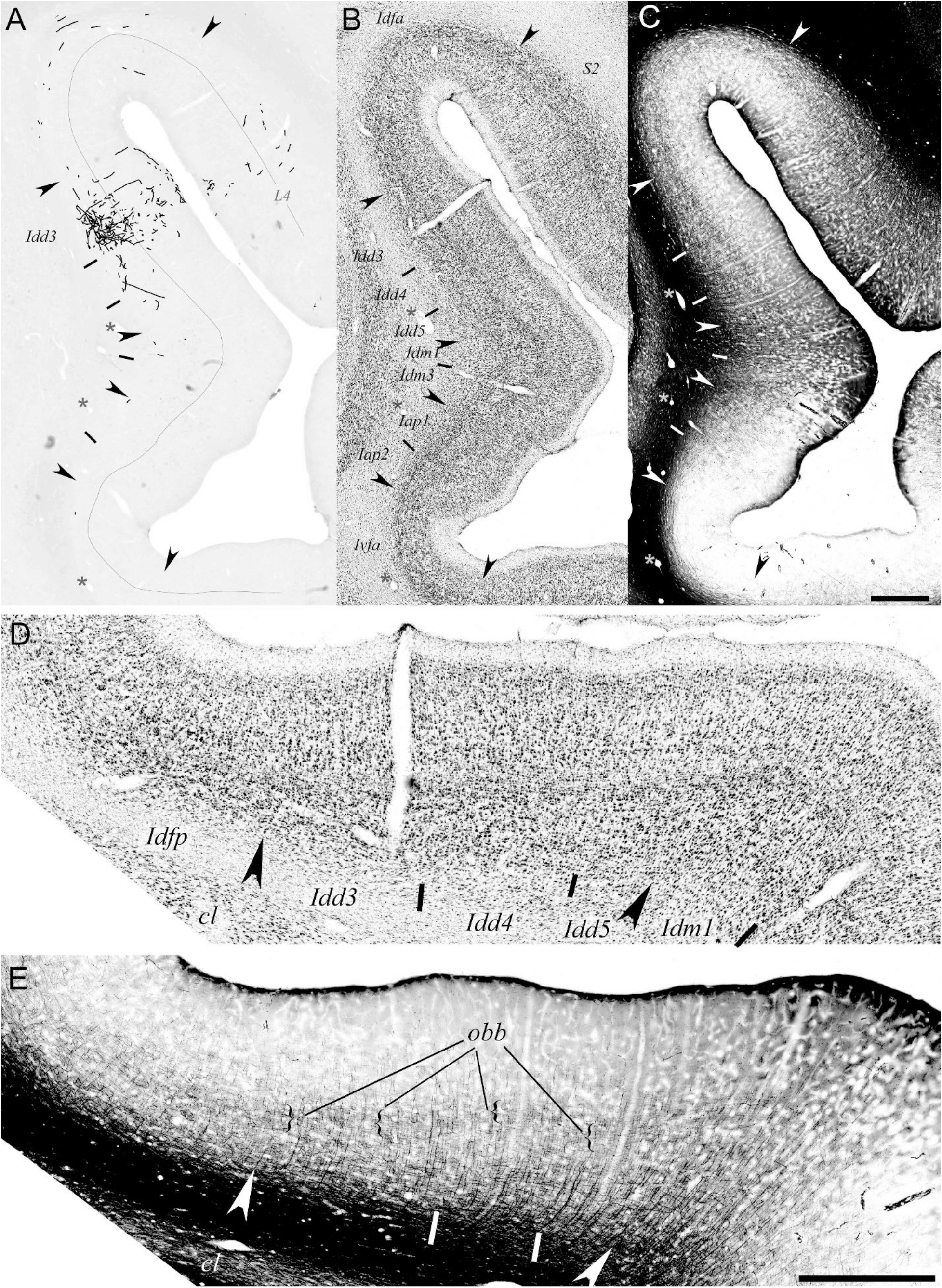
Composite showing the spatial overlap of patches of anterograde labeling with specific cyto- and myelo-architectonic subareas in the insular cortex in a macaque monkey (OM30) with an injection of biotin dextran amine (BDA) in the area 12m of the orbital prefrontal cortex (OPFC). **A.** Overlay of the charts of BDA-positive axon terminals on low-magnification photomicrographs of the corresponding BDA-stained sections of the insula. **B** and **C.** Overlay of the demarcation of, respectively, the cyto- and myelo-architectonic boundaries of the insula on low-magnification photomicrographs of Nissl- and Gallyas-stained sections directly adjacent to the BDA-stained section shown in panel A. All areas and subareas are noted. (See abbreviations list.) **D** and **E**. High-magnification view of the cyto- and myelo-architecture and boundaries in the dorsal half of the insula from panels B and C. In panel E, the outer band of Baillarger is indicated (obb). Scale = 1 mm.

**Figure 13.**
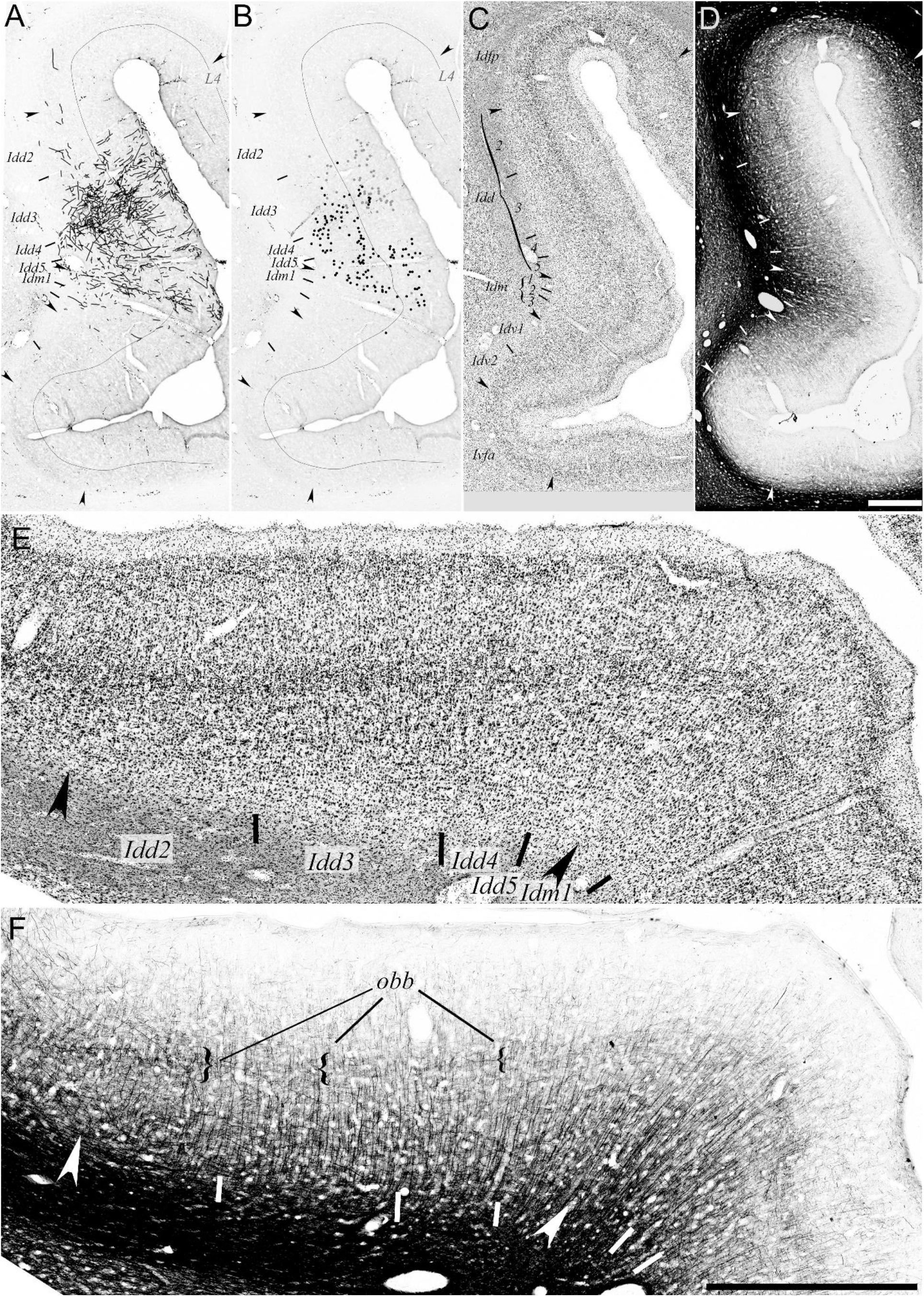
Composite showing the spatial overlap of patches of anterograde labeling with specific cyto- and myelo-architectonic subareas in the insular cortex in a macaque monkey (OM69) with an injection of lucifer yellow (LY) in the area 12r of the orbital prefrontal cortex (OPFC). **A-B.** Overlay of the charts of LY-positive axon terminals (A) and cell bodies (B) on low-magnification photomicrographs of the corresponding LY-stained sections of the insula. **C**-**D.** Overlay of the demarcation of the cyto- (C) and myelo- (D) architectonic boundaries of the insula on low-magnification photomicrographs of Nissl- and Gallyas-stained sections directly adjacent to the BDA-stained section shown in panel A. **E-F**. High-magnification view of the cyto- and myelo-architecture and boundaries in the middle of the insula from panels C and D. In panel F, the outer band of Baillarger is indicated (obb). (See abbreviations list.) Scale = 1 mm.

#### OM85

Labeling in OM85 occurred in a large middle portion of the insula, from Idfp to Ivfa, as well as in a small region within Ial (Fig. 3D). Figure 4A shows a representative low magnification photomicrograph of a coronal section from the middle portion of the insula, immunoreacted to visualize the anterograde and retrograde labeling produced by the injection of LY in area 24b’. Two conspicuous and sharply delimited patches of labeling are readily visible at this low magnification. Figure 4B shows the corresponding plot of individual retrogradely labeled perikarya and anterogradely labeled terminals. Figures 4C and 5 show the localization of cytoarchitectonic borders on an adjacent Nissl-stained section, at low and high magnifications, respectively. The same borders were transposed on the plot of anterograde and retrograde labeling (Fig. 4B).

From dorsal to ventral, we observed two regions with moderate anterograde labeling that corresponded to Idfp and Idd2, with clearly more labeling in Idd2 than in Idfp (Fig. 4B). The architectonic border between Idfp and Idd2 was characterized by a distinct vertical shift of the position of the slightly less discernable layer 4, a thickening and more collimated organization of layer 3 with barely discernible sub-layers 3a and b, and a broader layer 5 in Idd2 compared to Idfp (Fig. 4C and 5A). The next region of labeling corresponded to Idd3 and presented one of the most obvious and edifying cases of overlap between a tract-tracing labeling ‘patch’ and an architectonic sub-area (Fig. 4A-C and 5A). Idd3 contained a large group of darkly and moderately retrogradely labeled neurons in layer 3b and a dense mesh of anterogradely labeled fibers in layers 1 to 5. Architectonically, Idd3 was characterized by the presence of a thin but densely populated (‘darker’ in appearance) and very distinct layer 4 with particularly sharp delimitations (already distinct at low magnification) at the borders with Idd2 and Idd4, in which layer 4 was broader and less discernable (Fig. 4C and Fig. 5A). Idd3 had clearly separate and equally thick layers 3a and 3b, populated in this case with much ‘darker’ neurons than in Idd2. Layer 5 was slightly more compact and populated with smaller cells than in Idd2 and thinner than in Idd4. In contrast to Idd3, the moderate labeling in Idd4 was characterized by the presence of a thin ‘band’ of retrogradely labeled cells (contained within an equally thinner layer 3b), and distinctly less anterograde labeling, occurring mainly in outer and inner layers but not much in middle layers (Fig. 4B). Architectonically, in comparison to Idd3, Idd4 had a broader but less distinct layer 4, a broader layer 3 with a thin but poorly discernable layer 3b and larger layer 3a, and broader layers 5-6 (Fig. 4C and 5A). Idd5 had sparse anterograde labeling and no retrograde labeling (Fig. 4B). It was architectonically characterized by a thinner and less populated (sparser) layer 3 and larger cells in layer 5 (Fig. 4C and Fig. 5B).

Idm1 contained again a large population of retrogradely labeled neurons located this time both in layers 3 and 5, which differed from Idd3 in which labeled neurons occurred only in layer 3 (Fig. 4B). Anterogradely labeled fibers occurred across all layers. Idm1 was architectonically characterized by an outward shift in layers 3 to 5, a more collimated layer 3, and a broader layer 5 populated by smaller and more lanceolate neurons than in Idd5 (Fig. 4B and C). The ventral delimitation of the labeling in Idm1 coincided precisely with the architectonic border with Idm2 in which there was only sparse labeling (Fig. 4B). Idm2 differed from Idm1 by a thinner, sparser and less organized layer 3 and a distinct outward shift of layer 4 (Fig. 5C).

Ventral to Idm2, Idv1 and Idv2 contained dense and moderate anterograde and retrograde labeling, respectively (Fig. 4B). The anterograde labeling occurred in layers 1-3a and the labeled somas were in layer 3a and b in Idv1. The anterograde labeling in Idv2 was concentrated in layers 1, 3b and 5, and the cell bodies in layer 5, which contrasted with the layer distribution in Idv1. Architectonically, Idv1 had a thinner layer 5-6 and more homogeneous layer 3 (Fig. 5C). Idv2 had even thinner layers 5-6 than in Idv1, and was distinguished from Ivfp by a larger and small-celled layer 3 (Fig. 5C). Ivfp and Ivfa had dense and sparse anterograde labeling and no retrograde labeling. Ivfp had a thin, compact layer 5, thin layer but discernable layer 4, and thin layer 3 made of larger cells than in Idv2. Ivfa overall smaller cells than in Ivfp and no clearly discernable layer 4.

#### OM42

Anterograde labeling in OM42 occurred in a few dysgranular sub-areas in the middle portion of the insula, and more anteriorly in the rostral portion of Idm3, in Iap1, Ial, Iai, Iapm and Iam (Fig. 3E). The labeling filled the entire extent of Iapl and Iam, and a small fraction of Ial and Iai. Figures 6A-D show the plot of anterogradely labeled terminals overlaid on low magnification photomicrographs of the corresponding BDA-stained sections of the insula, at 4 levels passing through the Idd2 labeling patch (2500, 2000, 1500 and 1000 mm from the anterior commissure; 0.5-mm interval). Figures 6A’-D’ and 7 show the localization of cytoarchitectonic borders on, respectively, low and high magnification photomicrographs of Nissl-stained sections of the insula adjacent to the BDA-stained sections. (Figure 7 focuses on Idd2 and neighboring sub-areas.) The same borders were transposed on the BDA photomicrographs and plots in Figure 6A-D. Most noticeably, a heterogeneous but continuous patch of anterograde labeling formed an anteroposterior strip in Idd2 and, to some extent within Idfp. In addition to the patch in Idd2, anterograde labeling occurred in Idd5 across two consecutive sections (Fig. 6A and B), as well as in Idm1-3 in one section (Fig. 6D). The cytoarchitectonic boundary between Idfp and Idd2 was marked by a thickening layers 3 and 5, and a more diffuse layer 4 in Idd2. The outward shift of layer 4 observed in OM85 was not as pronounced in OM42 but still visible for instance in Fig. 7B and C. Conversely, Idd3 had equally thick layers 3a and 3b with sparser, larger and often darker neurons in layer 3b. Layer 5 was thinner and had smaller cells than in Idd2. Idd4 had a more diffuse layer 4 than in Idd3, with a less distinct separation from layers 3 and 5. As in OM85, layer 3a and b were also less separable in Idd4 than in Idd3.

#### OM27

The injection of BDA in area 11l in OM27 produced anterograde labeling in ta dorsal-posterior region and in a ventral-anterior region (Fig. 3F). The dorsal posterior labeling included labeling in Idfp, Idd1, 2, 3 and 4. The ventral anterior labeling included Idm2 and 3, Idv1 and 2, Iap1 and 2, and small portions of Iapl, Ial and Iai. Most labeled dysgranular sub-areas contained both anterograde and retrograde labeling. Agranular areas contained mainly anterograde labeling. Figures 8 to 11 focus on sections at four different ap levels (4500, 3000, 1500 and 0 mm from the anterior commissure; 1.5-mm interval) with labeling in both regions. Figures 8A-D show the plots of BDA-positive terminals overlaid on the low magnification photomicrographs of the corresponding BDA-reacted sections of the insula. Figures 8A’-D’ show the cytoarchitectonic boundaries on low magnification photomicrographs of adjacent Nissl-stained sections. Figures 9–11 show high magnification photomicrographs demonstrating the architectonic features of the subareas containing labeling.

Idfp contained a small patch of weak to moderate labeling, at the same anteroposterior levels where Idd1 contained moderate to dense labeling (Fig. 3F and 8A). Anterograde labeling occurred throughout a long segment within the rostrocaudal extent of Idd2, with however a noticeable gap, approximately in the middle of the segment, which coincided with an increase in labeling density in Idd1 (Fig. 3F and 8A-C). This local variation illustrates the complexity or non-triviality of the modular organization of the prefronto-insular connections. Figures 9A-C show high magnification photomicrographs of the dorsal portion of the Nissl-stained sections shown in Figure 8A’-C’. In all three sections, Idd1 differed from Idd2 by an overall thinner, denser and less columnar layer 3 with smaller cells than in Idd2, as well as a thinner and less populated layer 5. Idd2 showed sparser but larger cells with a slightly more collimated organization in its broader layer 3. The border between Idd2 and Idd3 was equally abrupt. Layer 3 in Idd3 was thinner with sparser but larger cells and a better distinction between layers 3a and 3b. The columnar organization seen in Idd2 was replaced with a more random organization in Idd3.

Figure 10 shows the cytoarchitectonic borders of Idd4, which contained a patch of anterograde labeling throughout its caudal two thirds (Fig. 8F). In comparison to Idd3, Idd4 had a thinner and more homogeneous layer 3 made of darkly stained pyramidal somas and a less distinct layer 4. The border between Idd4 and Idm1 was rather sharp, with a denser layer 3 made of smaller and more collimated cells in Idm1, an outward shift in layer and a markedly thicker layer 5 populated with smaller neurons, compared to Idd4.

In the mound area, Idm1 was completely devoid of labeling, conversely to Idm2, which contained dense labeling, with an abrupt dorsal delimitation that perfectly coincided with the cytoarchitectonic boundary between Idm1 and Idm2 (Fig. 8A-C). The boundary was marked by a subtle but definite fault line in the layer organization. The small and lanceolate cells in layers 2 and 3 in Idm1 were replaced with sparser, larger and rounder cells in Idm2. The discrete columnar organization was replaced with a random organization, in both layers 3 and 5. Idm3 had a varying density of anterograde labeling, from weak to moderate. However, the labeling was always distinct from the labeling in Idm2 and Idv1. The boundary with Idm2 was marked by a thinner layer 3 with even sparser neurons and a thinner homogeneous layer 5. Idv, as a whole, contained moderate to dense labeling with no clear distinction between Idv1 and Idv2, except perhaps for a progressive increase in density from Idv2, posteriorly, to Idv1, more anteriorly (Fig. 2F). The boundary between Idm3 and Idm1 was marked by a thickening of layer 3, with smaller and lanceolate cells, and less discernible layers 2 and 4. The boundary between Idv1 and Idv2 was marked by a sharp reduction in the thickness of layers 2, 3 and 5 (Fig. 11A-C). Finally, Ivfp had only sparse labeling at the border with Idv2 in a few sections. The architectonic border was marked by further thinning of all layers, and a sparser layer 3.

#### OM30

The anterograde labeling produced by the injection of BDA in area 12l was limited to a few patches (Fig. 3G). Figure 12 illustrates the sharp delimitation of one of these patches and its overlap with cyto- and myelo-architectonic boundaries of Idd3. Idd3 was characterized by a larger layer 3 than in Idfp, with distinct and equally thick layers 3a and b. Idd4 had a more homogenous layer 3 and a broader and also rather homogenous layer 5 (Fig. 12B and D). (Idd5 had a markedly sparser layer 3, compared to Idd4. Idm1 had broader layers 3 and 5 made of small, lanceolate cells arranged in narrow columns.) In the Gallyas stain, Idd3 had longer and coarser radial/vertical fibers than in Idfp and a diffuse outer band of Baillarger. Idd4 had even longer and coarser radial fibers and its outer band of Baillarger was sifted outwardly. (As in prior cases, the myelin stain density increased in Idd5 and the outer band of Baillarger was shifted inward and less distinct than in Idd4. Idm1 had very long, distinct and thin radial fibers.)

#### OM69

The injection of LY in OM69 produced anterograde and retrograde labeling in Idfp and Idfa, over a large region in the dorsal and to a lesser extent mound dysgranular areas, in Ivfa, Iapl, Iai and Iapm (Fig. 3H). Figure 13 illustrates the overlap of several adjacent patches of anterograde (Fig. 13A) and retrograde (Fig. 13B) labeling with cyto- and myelo-architectonic dorsal and mound dysgranular subareas. Indeed, a close comparison with adjacent Nissl- and Gallyas-stained sections revealed that the labeling was distributed across four distinct adjacent dysgranular subareas (Idd3, Idd4, Idd5 and Idm1) with a distinct pattern of anterograde and retrograde labeling in each subarea. From dorsal to ventral, Idd2 contained only sparse anterogradely labeled terminals, with a shape delimitation with Idd3 which contained conspicuous anterograde and retrograde labeling (Fig. 13A and B). The anterograde labeling in Idd3 was located mainly in layer 6, 5b and 3b, with sparser labeling in the other layers. The retrograde labeling was of moderate intensity (see Fig. 2 for reference) and located in both layer 5b and 3. Idd4 also contained anterograde and retrograde labeling. However, the anterograde labeling in Idd4 was slightly sparser than in Idd3 and the labeled somas occurred exclusively in layers 5a and 6. The anterograde labeling in Idd5 was similar to Idd4, but the retrograde labeling was located in layers 3, 5 and 6, and it was of stronger intensity than in Idd3. Finally, the labeling in Idm1 was characterized by a thin and almost continuous band of labeled somas from layer 6 to 3b, and a bundle of terminals spanning across all layers with the exception of layer 3a. Conversely, Idm2 contained only very sparse anterograde labeling and no retrograde labeling.

The cytoarchitecture in OM69 was not as obvious as in the other cases due to noise created by a contamination of the tissue with stained blood cells that did not wash optimally during the perfusion. Nevertheless, the main modular cytoarchitectonic features could be distinguished (Fig. 12C and E) and matched with the myeloarchitectonic features (Fig. 13D and F). The borders between the distinct dorsal dysgranular subareas were marked by changes in the thickness and demarcation of granular layer 4, as well as some variations in the cellular organization of layer 3 and to some extent 5. Like in the other cases, layer 4 was thin in Idd2, thicker and more distinct in Idd3, very diffuse in Idd4, and broader and slightly more recognizable in Idd5. The border between Idd2 and Idd3 was also marked with a sudden decrease is cell density in layer 3 as well as a thinner but large-celled layer 5 in Idd3, compared to Idd2. In addition of the diffuse layer 4, Idd4 had a thinner and more homogeneous layer 3 than in Idd3. The border between Idd4 and Idd5 was subtly marked by an outward shift in layer 5, a thinner layer 3 and broader layer 5 with smaller cells than in Idd4. Idm1 had overall smaller, more lanceolate and collimated cells, with a broader layer 5, similar to other cases. In parallel with the changes in the cellular appearance of layer 4, the outer band of Baillarger was very distinct in Idd2, weak in Idd3 and Idd4, and not distinct in Idd5 (Fig. 13F). Idd2 had a coarse mesh of stained fibers in layers 5 and 6 which differed from the coarse and more radial (vertical) fibers in Idd3. The difference between Idd3 and Idd4 was tenuous with only slightly less horizontal fibers crossing the radial fibers in Idd4. Idd5 had longer and straighter radial fibers than in Idd4. Idm1 was very distinct, with long radial fibers crossing the weak band of Baillarger into layer 3.

## Discussion

The present neuroanatomical report demonstrates the close spatial relationship between the localization of architectonic boundaries and the delimitation of discrete regions of anterograde and/or retrograde labeling in the insular cortex in the macaque monkey. The architecto-hodological overlap validates the recent architectonic parcellation of the classical granular, dysgranular and agranular sectors of the macaque insula into smaller and rather sharply delimited areas and subareas (Evrard *et al.* 2014), which is consistent with modern reports demonstrating a similar overlap in other cortical lobes (*e.g.*, Lewis and Van Essen 2000; Kaas 2002; Price 2007; Qi *et al.* 2008). Conversely, the overlap definitely refutes a former conclusion that “connectivity patterns” have “negligible” value for the determination of cytoarchitectonic borders in the primate insula (Gallay *et al.* 2012).

Although quite straightforward in appearance, the present finding represents a major progress and novelty in the examination of the primate insula. It will become of paramount importance for models of insular processing. Even if the ‘overlap’ may not be the rule for all projections to and from the insula (see below), it provides for the first time a clear structural support for the notion that the primate insulo-insular processing occurs within a refined modular *Bauplan,* in which each dysgranular subarea likely integrates interoception with functionally distinct polymodal and limbic afferent activities (Evrard and Craig 2015; Evrard 2018).

### Comparison with prior tract-tracing studies in macaques

Numerous prior tract-tracing studies reported labeling in the insula (*e.g.*, Burton and Jones 1976; Jones and Burton 1976; Pandya *et al.* 1981; Mesulam and Mufson 1982; Mufson and Mesulam 1982; Cavada *et al.* 1984; Matelli *et al.* 1986; Selemon and Goldman-Rakic 1988; Preuss and Goldman-Rakic 1989; Deacon 1992; Carmichael and Price 1995; Chikama *et al.* 1997; Cipolloni and Pandya 1999; Stefanacci and Amaral 2000; Lavenex *et al.* 2002; Fudge *et al.* 2005; Borra *et al.* 2008). The labeling never entirely ‘filled’ one of the three classical cytoarchitectonic sectors. It was instead heterogeneous and a close examination of figures from these studies readily reveals a more complex organization that often included ‘patches’ and ‘stripes’ fitting with the present finding.

One of the most edifying examples comes from a study by Selemon and Goldman-Rakic (1988). In this study, large injections of horseradish peroxidase in prefrontal cortex and tritiated amino acid (TAA) in posterior parietal cortex produced sharply delimited and overlapping patches of anterograde labeling in an insular region that could correspond to one of the architectonic subareas of our dorsal dysgranular area (Fig. 14A). In another example, Burton and colleagues (1995) showed “stripes” of retrograde labeling that were continuous across almost several mm. They located these stripes in the granular insula (“Ig” in Fig. 14B) but they could very well correspond to the first and third (Idd1 and 3) or second and fourth (Idd2 and 4) subareas of our dorsal dysgranular area. This example directly supports the continuous anteroposterior patches of labeling reported in the present study. In a final example, large injections of TAA in the frontal operculum, posterior parietal cortex or prefrontal cortex also produced rather heterogeneous anterograde labeling in the insula (Mufson and Mesulam 1982). The flat map illustrations of this labeling (Fig. 14C) clearly denoted small anteroposterior stripes that were similar to the stripes reported in the present contribution (Fig. 3).

**Figure 14.**
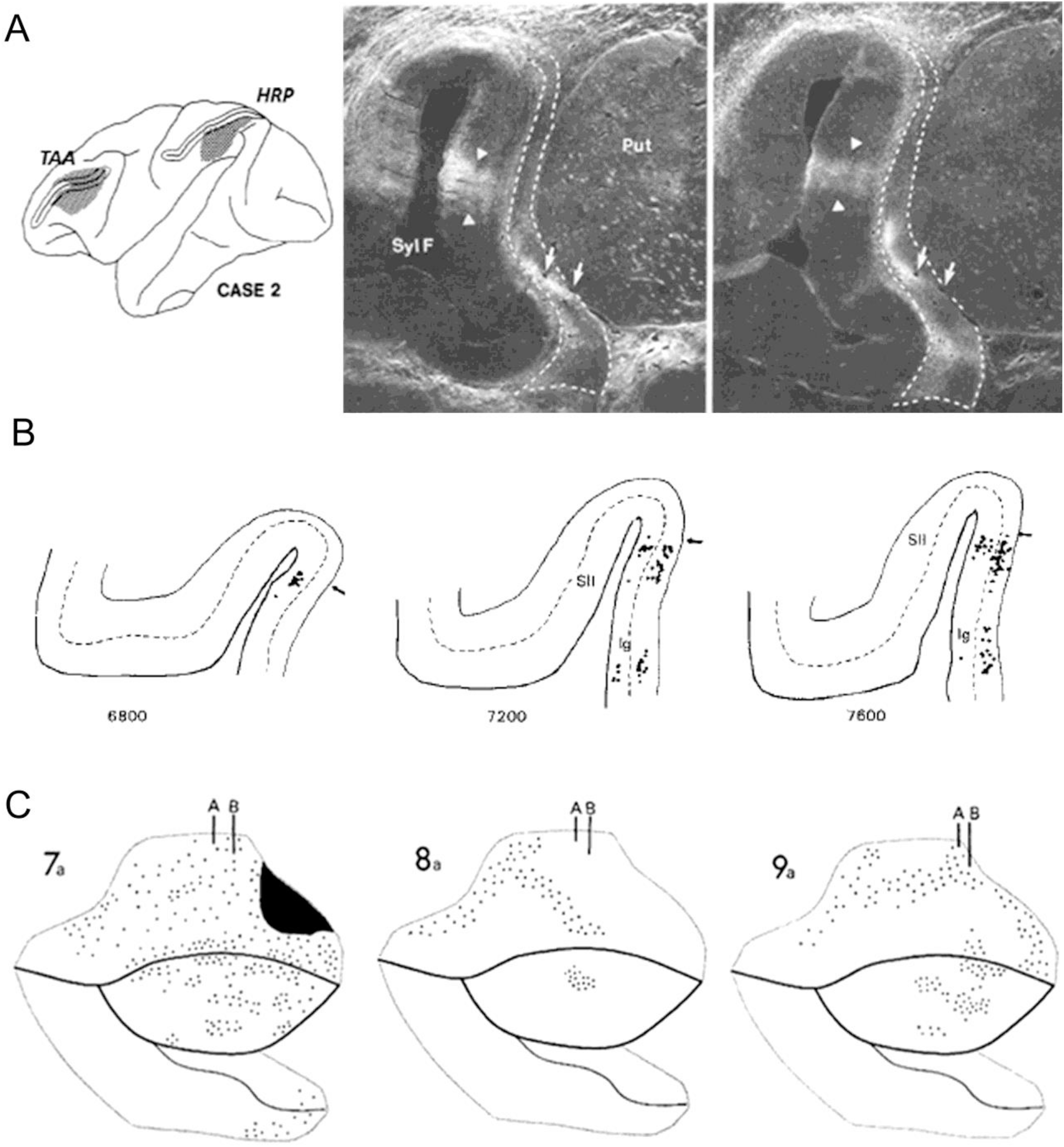
**A.** Coronal photomicrographs showing patches of tritiated amino acid (TAA, left) and horseradish peroxidase (HRP, right) anterograde labeling in the insula with injections in the prefrontal and posterior parietal cortices (Adapted with permission from Selemon & Goldman-Rakic, 1988). **B.** Distribution of DY-labeled cells on selected coronal sections through the dorsal dysgranular insula after injections in the distal toe region of area 3b. Each dot represents the location of one retrogradely labeled cell (Adapted with permission from Burton et al., 1995). **C.** Flat map representation of “stripes” of retrograde labeling in the insula in 3 monkeys with injection of HRP in distinct parts of the brain. The injections were made in the frontoparietal operculum (7a), posterior parietal cortex (8a) and lateral prefrontal cortex (9a) (Adapted with permission from Mesulam and Mufson, 1982).

In their seminal tract-tracing studies of the cortico-cortical connections of the macaque insula, Mesulam and Mufson (1982b) divided the insula into two vast hodological territories (Fig. 1F). They proposed an anteroventral ‘limbic’ territory including the agranular insula and the rostral half of the dysgranular insula separated by a gradual transition from a posterodorsal ‘auditory-somesthetic-skeletomotor’ territory including roughly the caudal half of the dysgranular insula and the granular insula. While this general anteroposterior progression will likely remain valid and valuable (Cerliani *et al.* 2010; Cerliani *et al.* 2012), the present finding recommends a thorough reexamination of all existing and viable tract-tracing material with labeling in the insula. The examples above, and in particular the case of dual anterograde labeling in one single discrete region with two very distinct injections (Fig. 14A), readily announce that such reexamination would definitely enlighten our understanding of both the anatomical and functional organization of the primate insula.

### Comparison with other species

We already reviewed in our prior contribution the major organizational and topographical differences between the monkey, cat and rat insulae, including for instance the absence of a distinct interoceptive spinothalamocortical pathway (see *Discussion* in Evrard *et al.* 2014). Tract-tracing studies of the rat insula demonstrated optimal overlaps between architectonics and hodology (*e.g.*, Allen *et al.* 1991; Shi and Cassell 1998; Jasmin *et al.* 2004). However, this overlap was limited to the three-five classical architectonic areas of the rat insula and did not reveal thinner ‘modular’ parcellation. In addition, the projections from the putative analog of the monkey prefrontal cortex targeted mainly the agranular area of the insula (Allen *et al.* 1991; Gabbott *et al.* 2003), whereas in macaque the prefrontal cortex projects in a very heterogeneous manner chiefly to the dysgranular areas and to a lesser extent to the agranular areas. Although an overall involvement in homeostasis can still offer points of comparison across species and lead to the enunciation of general analog principles, the modular and hodological organization of the macaque insula is, by far, more elaborate than that of the rodent, and justifies using caution when comparing this structure between primate and non-primate species. The addition of areas and subareas in the macaque insula likely reflects major evolutionary adaptations and additions of new functions, as already proposed for other systems (Kaas 2002; Kaas 2004).

The existence of multiple sharply-delimited insular areas is common to our work and these of Rose (1928), Brockhaus (1942), and Kurth *et al.* (2010) in humans (Fig. 1) (See *Discussion* in Evrard *et al.* 2014). Additional architectonic examinations of the human insula are now needed to analyze (1) whether the three dysgranular areas posterior to the central sulcus (Kurth, Eickhoff, *et al.* 2010) contain finer subdivisions, (2) whether these subdivisions continue passed the central sulcus into the small gyri (which have not yet been mapped) or are limited to the posterior long gyri, (3) whether these putative posterior subdivisions are aligned parallel to the dorsal fundus or to the axis of the long gyri, (4) whether the anterior small gyri and the ‘frontoinsula’ are architectonically divided into more areas than in macaques as suggested by their recent disproportional expansion of the anterior insula in humans (Bauernfeind *et al.* 2013), and, finally, (5) whether the von Economo neurons which seem to occur in only one architectonic area in macaques (Horn *et al.* 2017) do indeed occur in more than one area in humans, as suggested by our preliminary study (Horn and Evrard 2018; Horn and Evrard, *in preparation*). Recent functional studies in humans suggested the presence of several distinct anatomical and functional domains within the human insula (Kurth, Zilles, *et al.* 2010; Deen *et al.* 2011; Kelly *et al.* 2012). Continuous efforts to obtain more accurate functional maps of the insula using high-resolution fMRI and to compare these with maps obtained in the macaque monkey may help decipher fixed homologies from species-specific innovations.

### Cortical patterns and insular map

The cerebral cortex contains a series of maps that reflect the topography and superposition or separation of modalities within distinct sensory systems. These maps tend to represent peripheral changes and discontinuities in ‘primary’ sensory cortices and evolve towards more ‘abstract’ polymodal maps when progressing towards higher association areas. The organization of these maps, regardless of the level of integration, varies greatly, depending on the sensory modality, underlying connection patterns, species, etc. (Swindale 1990; Horton and Adams 2005; Kaas 2012). In the present study, the fine dysgranular sub-areas in the insular map do not refer to highly regular and periodic columnar patterns such as these observed in the visual or somatosensory cortex in some species (Mountcastle *et al.* 1957; Hendrickson 1985; Purves *et al.* 1992; Mountcastle 1997). Although we previously defined our dysgranular subareas as “modules” (Evrard et al., 2014), our subareas are unlikely to represent receptive field discontinuity such as these observed in the “modules” put forward by Kaas in sensory areas across modalities and species (Kaas 2012). The insular “patches” and “stripes” in the present study bear resemblance with prior patterns of periodic intrinsic cortico-cortical projections (Levitt *et al.* 1993; Lund *et al.* 1993) and interdigitated extrinsic projections to the frontal association cortex (Goldman-Rakic and Schwartz 1982). However, these patterns occurred within single areas without evidence of overlap with finer sub-areal architectonic parcellations. While repeatability and complementary interdigitating are not excluded, one of the large prefrontal injections labeled only one small region within a subarea without any evidence of repetition (OM30), while one of the small injections labeled multiple adjacent patches without leaving much room for an interdigitated pattern (OM27). Rather, as detailed in our earlier model, we suggest that the fine horizontal ‘stripe-like’ partition of the insula proper reflects the occurrence of a serial processing stream where interoception is integrated with multi-modal activities in order to provide a rather unified representation of interoceptive, exteroceptive and teloreceptive activities of the organism (Evrard and Craig 2015; Evrard 2018; Evrard 2019).

The dorsal fundus of the insular cortex is the terminus of a primate-specific spino-thalamocortical pathway encoding the physiological state of the body and constitutes the “primary interoceptive cortex” (I1) (Craig 2002). In our working model of the insula, each horizontal dysgranular subarea receives inputs from I1 and integrates them with afferent activities from selective sets of cortical and subcortical regions (Evrard and Craig 2015; Evrard 2018). This dorso-ventral homotopic insulo-insular communication is supported by recent tract-tracing in our lab, showing that one small injection of tracer in a restricted portion of I1 produces labeling in equally restricted portions of the dysgranular subareas at the same anteroposterior levels (Welzel and Evrard, *unpublished observations*; see also). This trans-areal process is consistent with the idea that the representation of core affect and sentient feelings in the anterior insula derives from the integration of ongoing environmental signals with interoceptive and homeostatic activities in the posterior and middle insula (Evrard and Craig 2015).

## Acknowledgments

This work was supported by the Werner Reichardt Centre for Integrative Neuroscience (CIN) at the Eberhard Karls University of Tübingen (CIN is an Excellence Cluster funded by the Deutsche Forschungsgemeinschaft [DFG] within the framework of the Excellence Initiative EXC 307) and by the Max Planck Society. Thanks to Prof. Dr. Joseph L. Price for allowing us to use his valuable tract-tracing collection and Prof. Dr. Arthur Lowey for providing generous access to his lab and microscopes, both at the Washington University of Saint-Louis, MO, USA.

## Abbreviations

*BDA*: Biotinylated dextran amine
*CTb*: Cholera toxin subunit B
*FR*: Fluororuby
*Iai*: intermediate agranular area of the insula
*Ial*: lateral agranular area of the insula
*Iam*: medial agranular area of the insula
*Iap1-2*: posterior agranular area of the insula, subareas 1 and 2
*Iapl*: posterior lateral agranular area of the insula
*Iapm*: posterior medial agranular area of the insula
*Idd1-5*: dorsal dysgranular area of the insula, subareas 1 to 5
*Idfa*: anterior (granular) area of the dorsal fundus of the insula
*Idfp*: posterior (granular) area of the dorsal fundus of the insula
*Idm1-3*: mound dysgranular area of the insula, subareas 1 to 3
*Idv1-2*: ventral dysgranular area of the insula, subareas 1 and 2
*ILS*: Inferior limiting sulcus of the insula
*Ivfa*: anterior (agranular) area of the ventral fundus of the insula
*Ivfp*: posterior (dysgranular) area of the ventral fundus of the insula
*LY*: Lucifer yellow
*SLS*: superior limiting sulcus of the insula

